# P-Rex2 exhibits unique structural features and regulatory mechanisms distinct from the closely related RhoGEF P-Rex1

**DOI:** 10.64898/2026.01.05.697791

**Authors:** Lauren K. Anderson, Rohan Marde, Grace Muma, Veda Nayak, Chi Phan, Sheng Li, Jennifer N. Cash

## Abstract

Rho guanine-nucleotide exchange factors (RhoGEFs) activate small GTPases to drive cytoskeletal rearrangement, cell motility, and proliferation. The phosphatidylinositol-3,4,5-trisphosphate (PIP_3_)-dependent Rac exchanger (P-Rex) subfamily of RhoGEFs includes P-Rex1 and P-Rex2 which, when misregulated, contribute to cancer progression and metastasis. P-Rex activity is controlled by accessory domains that maintain the protein in a cytosolic, autoinhibited state until activated by the lipid PIP_3_ and G protein βγ subunits. While P-Rex1 autoinhibition has been structurally and biochemically characterized, P-Rex2 has remained largely unexplored. Furthermore, despite high sequence similarity and domain conservation, P-Rex homologs differ in substrate specificity and regulatory interactions, and the molecular basis for these divergences is unknown. Here, we have taken an integrative structural biology approach to investigate these gaps. Using cryo-EM, we determined the first structure of full-length P-Rex2 to moderate resolution, revealing that, while the overall structure closely resembles that of P-Rex1, there is a substantial repositioning of the N-terminal module relative to the C-terminal core. This may play a key role in precluding the intramolecular interactions between the N- and C-terminal domains that are observed in autoinhibited P-Rex1. Hydrogen-deuterium exchange mass spectrometry revealed that, unlike P-Rex1, P-Rex2 dynamics are unaffected by IP_4_, the headgroup of PIP_3_. SEC-SAXS data support that the N-terminal module itself is less dynamic, and biochemical assays show that P-Rex2 may be more tightly regulated by autoinhibition, likely through a mechanism different from P-Rex1. These findings uncover unique features in the molecular mechanisms of P-Rex2 regulation.

## Introduction

Rho guanine-nucleotide exchange factors (RhoGEFs) accelerate the exchange of GDP for GTP on small Rho GTPases like Rac and Cdc42 to direct actin cytoskeleton rearrangement^1,2^. This activity allows them to drive cell migration and invasion which, when misregulated, can contribute to various diseases including cancer, resulting in metastasis^3^. Members of the Dbl family of RhoGEFs, which contains over 70 members, share a conserved Dbl homology (DH) catalytic domain followed by a pleckstrin homology (PH) domain that together form the catalytic core^4^. Although the DH/PH tandem is well characterized structurally^5–7^, the variety of accessory domains that accompany the catalytic core across and within different RhoGEFs introduces complicated, and comparatively vastly understudied, regulatory^*^ controls on GEF activity. These include regulation through autoinhibition, membrane localization, and protein-protein interactions^1,8^. As a result, the complex molecular mechanisms governing the activity of each of these multidomain proteins are poorly understood.

The phosphatidylinositol (3,4,5)-trisphosphate (PIP_3_)-dependent Rac exchanger (P-Rex) subfamily of Dbl RhoGEFs includes two members, P-Rex1 and P-Rex2. P-Rex1 is abundantly expressed in leukocytes and neurons whereas P-Rex2 is strongly expressed in brain, lung and liver cells^9–12^. *PREX2* additionally gives rise to a splice variant, P-Rex2b, which is mainly expressed in heart tissue and retains all identifiable domains except the IP4P domain which is truncated and ends with a short alternative sequence^13^. Misregulation of P-Rex family proteins is associated with varied pathologies. While P-Rex1-driven diseases, like prostate and breast cancers, are linked to overexpression^14–16^, P-Rex2-driven diseases are propelled by mutation. For example, more than 23% of human hepatocellular carcinoma and 38% of squamous cell carcinoma samples harbor non-silent somatic mutations in P-Rex2^17,18^. Additionally, mutations in P-Rex2 can lead to metastatic melanoma^19^ and pancreatic cancers^20^ or disrupt PI3K signaling pathways to drive diabetes and insulin resistance^21^. In contrast, single and double knock-out of *PREX1* and *PREX2* genes in mouse models produces healthy, viable mice with very mild neutrophilia and a characteristic pigmentation phenotype^12,22^. Collectively, these data suggest that P-Rex is an excellent therapeutic target. Thus, characterizing the molecular mechanisms involved in P-Rex1 and P-Rex2 regulation and defining the differences between the two are important for guiding efforts to rationally design isoform-specific P-Rex modulators, both for use as tools in understanding RhoGEF regulation and for potential treatment of diseases.

P-Rex1 and P-Rex2 show 56% sequence identity and 68% sequence similarity overall, and both proteins share a conserved domain architecture consisting of an N-terminal DH/PH tandem followed by two Dishevelled, Egl-10 and Pleckstrin (DEP) domains, two Postsynaptic density protein 95, Discs large protein, and Zonula occludens-1 (PDZ) domains, and a C-terminal inositol polyphosphate-4-phosphatase-like (IP4P) domain (Fig. 1A). The P-Rex1 N-terminus forms a multi-domain module composed of the DH, PH, and DEP1 domains which is flexibly tethered to the core (Fig. 1B). The DH domain, which engages GTPases directly, is highly conserved as are the other domains within the N-terminal module (Fig. 1A). P-Rex1 also contains a C-terminal “core” composed of the tightly associated DEP2, PDZ1, PDZ2, and IP4P domains which exhibit the highest sequence divergence between the two proteins (Fig. 1A&B). Furthermore, a flexibly tethered subdomain of ∼250 residues extends out from the IP4P domain and forms a 4-helix bundle (4HB).

**Figure 1.**
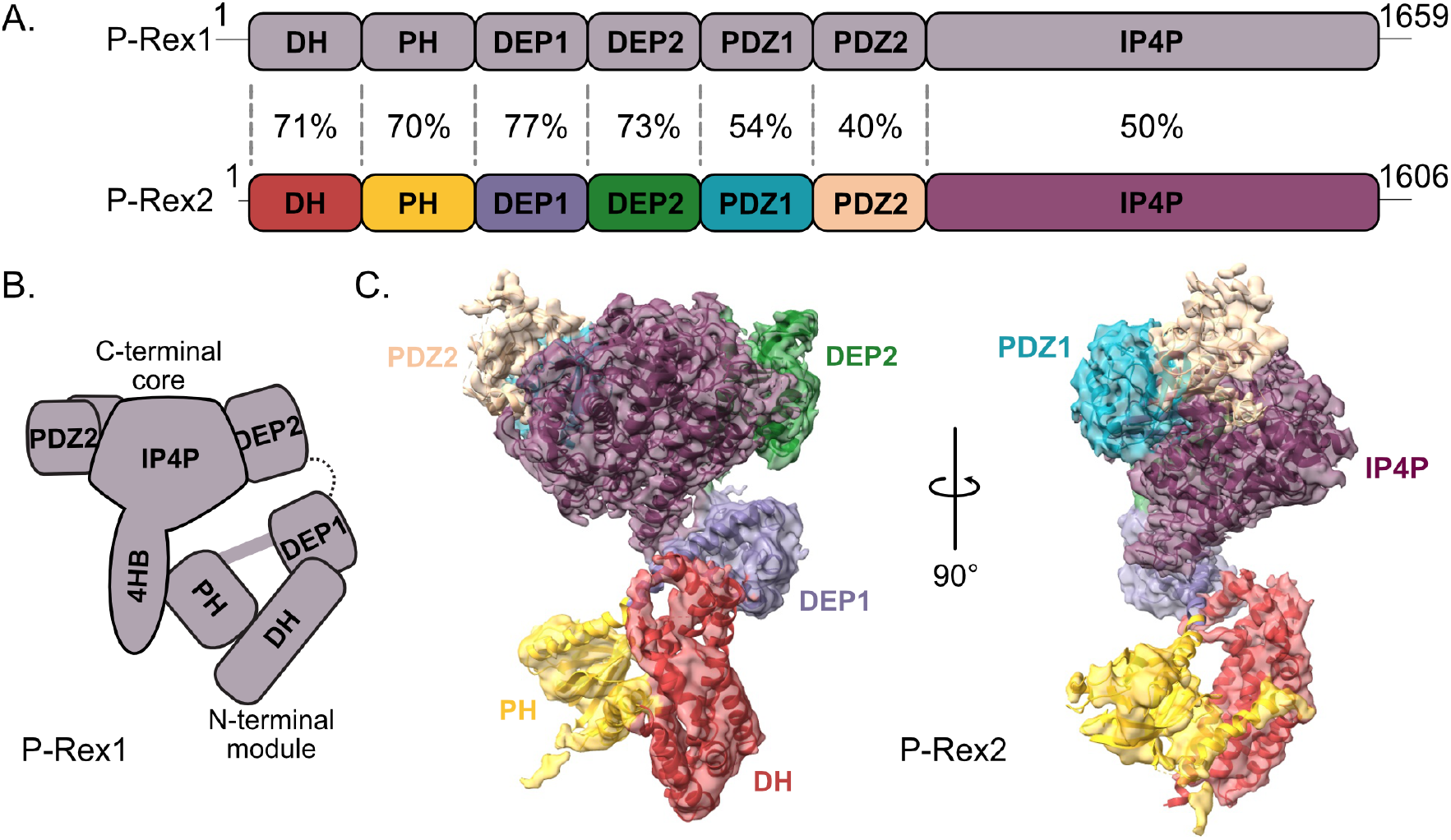
P-Rex2 exhibits a structure that is overall conserved with P-Rex1. A) Domain layout of P-Rex1 in gray and P-Rex2 colored by domain. The sequence identities between domains of the isoforms are labeled. B) Schematic of the domain organization of P-Rex1. C) Cryo-EM reconstruction of P-Rex2 (composite map) colored by domain with atomic model superimposed. The IP4P subdomain was not resolved in P-Rex2, unlike in P-Rex1.

Both enzymes are maintained in a cytosolic, autoinhibited state with limited basal activity until they are recruited to the cell membrane and synergistically activated by the lipid PIP_3_ and the Gβγ subunits of heterotrimeric G proteins^9,13,23^. Structural studies of P-Rex1 have begun to clarify how accessory domains facilitate autoinhibition and how Gβγ and PIP_3_ relieve this^5,24–29^. A two-pronged autoinhibitory mechanism is mediated by interdomain contacts that restrict the DH domain from interacting with GTPases^24,25^. One interaction is formed by the DEP1 domain docking against the DH domain through a hydrophobic pocket of residues to stabilize a compact, low activity state^27^ (Fig. 1B). A second interaction is mediated through the PH domain engaging the 4HB of the IP4P^24,25^. PIP_3_ binding to the PH domain disrupts the PH–IP4P contact— an effect that the soluble headgroup of PIP_3_, IP_4_, does not reproduce^25^. In fact, IP_4_ actually stabilizes the PH–IP4P autoinhibitory conformation. Gβγ, which binds to the surface formed by the DEP2, PDZ1, PDZ2 and IP4P domains, likely contributes to activation by promoting membrane localization, but this has not been conclusively determined^26^. Together, PIP_3_ and Gβγ bind to synergistically activate P-Rex1 and fully relieve autoinhibition. Despite the wealth of structural information available on P-Rex1^5,24–27,30^, there is only a single reported structure of a piece of P-Rex2: the isolated PH domain^31^.

While the close sequence similarity and shared domain architecture suggest that P-Rex2 adopts a structure similar to P-Rex1, there are notable differences between the two in substrate specificity and regulation^8,32^. For example, P-Rex1 can activate all Rac-like and some Cdc42-like GTPases *in vitro*, but only Rac-like GTPase activation has been observed *in vivo*^6,9,32,33^. On the other hand, P-Rex2 specificity *in vitro* seems to be limited to Rac-like GTPases^23,34^. Additionally, the tumor suppressor protein phosphatase and tensin homolog (PTEN) has been shown to form a co-inhibitory interaction with P-Rex2, but not P-Rex1^10,35,36^. Furthermore, P-Rex1 contains a membrane localization element in the β3/β4 loop of its PH domain that is missing in P-Rex2. When this element is swapped into the corresponding loop of P-Rex2, it enhances membrane localization of P-Rex2^5^. Collectively, this evidence suggests that there may be differences in the molecular features of P-Rex1 and P-Rex2 that lead to these unique properties. In support of this, cross-linking mass spectrometry data on P-Rex2^36^ do not entirely correlate with the autoinhibited structure of P-Rex1, as P-Rex1 homologs of some cross-linked P-Rex2 residues are too far apart— much beyond the length of the crosslinker utilized (Fig. S1A). These data point to potential structural differences between the two proteins that have not yet been described.

Here, we investigated the structure and autoinhibitory mechanisms of P-Rex2 using an integrative structural biology approach. We determined a moderate-resolution cryo-electron microscopy (cryo-EM) structure of full-length P-Rex2 which showed that, although the N-terminal module and C-terminal core each individually exhibit a similar domain organization as compared to P-Rex1, the N-terminal module itself is positioned proximally to the core, with the DEP1 domain contacting the base of the IP4P domain. Furthermore, hydrogen-deuterium exchange mass spectrometry (HDX-MS) experiments show that, unlike P-Rex1, binding of IP_4_ has no effect on the overall conformation of P-Rex2. Size exclusion chromatography coupled with small-angle X-ray scattering (SEC-SAXS) shows that although DEP1 promotes a compact overall architecture of the N-terminal module, it does not change the flexibility of the DH/PH tandem as previously described in P-Rex1. Despite these differences, mutagenesis and biochemical assays demonstrate that the DEP1 domain remains essential for autoinhibition by engaging the DH domain similarly to P-Rex1. This work reveals unique structural features that distinguish P-

Rex2 from P-Rex1, providing insights into key molecular details that may confer regulatory differences between the two.

## Results

### The overall structure of P-Rex2 resembles that of P-Rex1 with differences in the positioning of the N-terminal module

To begin to understand the molecular differences in P-Rex2 as compared to P-Rex1, we determined the structure of full-length (FL) P-Rex2 using cryo-electron microscopy (cryo-EM) single-particle analysis (Fig. 1C, S2-S4, & Table 1). Similar to P-Rex1, P-Rex2 exhibits a strong preferred orientation on grids (Fig. S2), necessitating the collection of supplemental data at a 35° tilt to provide additional views of the particle. Still, this was not enough to completely overcome the preferred orientation problem, and this coupled with inherent flexibility within P-Rex2 led to anisotropy within our maps. The overall domain organization of P-Rex2 resembles that of P-Rex1, composed of an N-terminal catalytic module (DH/PH-DEP1) and a C-terminal regulatory core (DEP2-IP4P) (Fig. 1C). The core was resolved to higher resolution overall, exhibiting a global resolution of 2.9 Å according to cryoSPARC (Fig. S2D-G), while the N-terminal module was resolved to a lower global resolution of 4.4 Å after focused refinement (Fig. S2H-J). Through an alternative data processing pipeline, we were also able to generate an additional map representing the whole particle at a global resolution of 3.4 Å (Fig. S2K-M).

**Table 1.** Cryo-EM Data Collection, Processing, and Modeling Statistics.

| Data Processing | Composite Map | Consensus Map | N-terminal Local Refinement | Whole Particle Local Refinement |
| --- | --- | --- | --- | --- |
| EMDB accession code | EMD-74550 | EMD-74547 | EMD-74548 | EMD-74549 |
| Symmetry imposed | C1 | C1 | C1 | C1 |
| Particles in final map | - | 215,042 | 101,178 | 128,298 |
| Global Resolution (Å, at 0.143) | - | 2.9 | 4.4 | 3.4 |
| 3DFSC sphericity | - | 0.734 | 0.872 | 0.83 |
| Map sharpening B factor (Å<sup>2</sup>) | - | 79.8 | 161.7 | 76.3 |

| Data Collection & Pre-Processing | Untilted | Tilted |
| --- | --- | --- |
| EMPIAR accession code | XXXX |  |
| Micrographs (collected) | 12,070 | 7,722 |
| Electron dose (e<sup>-</sup>/Å<sup>2</sup>) | 50 | 50 |
| Frames per movie | 1143 | 1206 |
| Voltage (kV) | 300 | 300 |
| Stage tilt (°) | 0 | 35 |
| Pixel size (Å) | 0.7296 | 0.7296 |
| Defocus range (µm) | 0.7-2.2 | 0.7-2.2 |
| Micrographs (used) | 8133 | 6959 |
| Extracted particles | 2,184,088 | 2,137,912 |
| Final particles after 2D classification | 1,390,781 | 1,585,976 |

| Atomic modeling |  |
| --- | --- |
| PDB accession code | 9ZQ7 |
| Model Composition |  |
| Chains | 1 |
| Non-hydrogen atoms | 8320 |
| Hydrogens | 7704 |
| Protein residues | 1035 |
| Water | 0 |
| Ligands | 0 |
| B factors (Å<sup>2</sup>, min/max/mean) |  |
| Protein | 30/524/132 |
| Ligand | 0 |
| Water | 0 |
| R.m.s. deviations |  |
| Bond lengths (Å) | 0.003 |
| Bond angles (°) | 0.477 |
| Validation |  |
| MolProbity score | 1.63 |
| Clashscore | 6.97 |
| Rotamer outliers (%) | 1.53 |
| CaBLAM outliers (%) | 0.86 |
| Ramachandran plot (%) |  |
| Favored | 97.5 |
| Allowed | 2.4 |
| Outliers | 0.1 |
| Model vs. Data* |  |
| CC mask | 0.58 |
| CC box | 0.7 |
| CC peaks | 0.44 |
| CC volume | 0.6 |
| Map-to-model FSC (Å, at 0.5)* | 3.4 |
| EMRinger score* | 1.87 |
| Q-score* | 0.37 |

\*Composite map

Like in autoinhibited P-Rex1, the P-Rex2 DH domain is connected to the PH domain through a bent helix, while the PH domain is connected to the DEP1 domain through a more linear, continuous helix, and the DEP1 domains docks onto the end of the DH helical bundle (Fig. 1C, 2 & S4B-D). The C-terminal core of P-Rex2 is composed of the second DEP domain, both PDZ domains and the IP4P domain where the IP4P domain is the central component and is decorated with the other accessory domains at peripheral ends. A β-strand connects DEP2 to PDZ1 and is part of a large β-sheet that twists through the center of the IP4P domain from one side to the opposite. The domain organization within each the N-terminal module and C-terminal core is conserved between P-Rex1 and P-Rex2, exhibiting an RMSD Cα of 1.4 Å each (Fig. 2). However, the P-Rex2 IP4P subdomain, observed in the autoinhibited P-Rex1 structures, is not resolved in any of our 2D class averages or 3D reconstructions.

**Figure 2.**
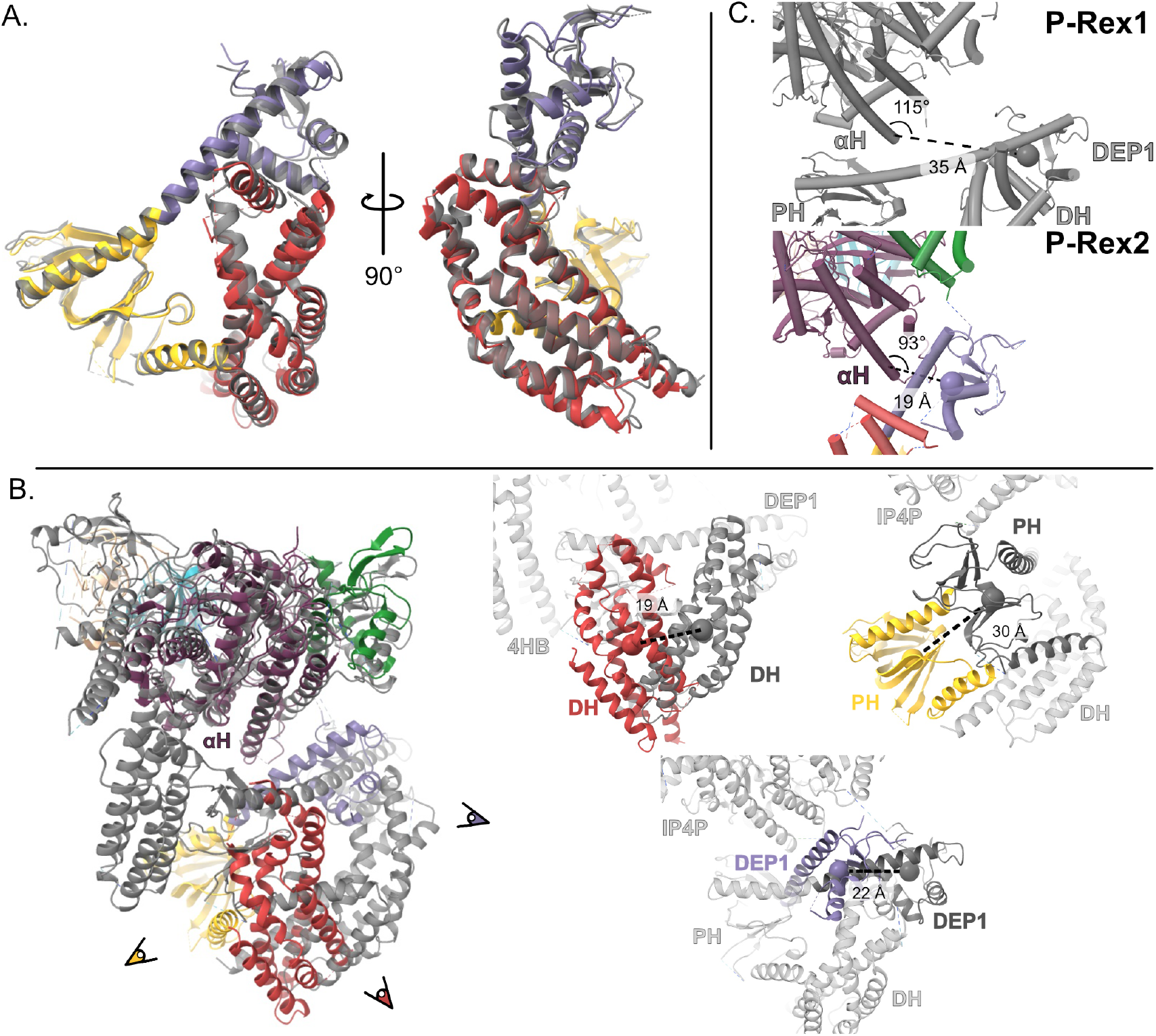
P-Rex2 shows a relative shift in the position of the N-terminal module as compared to P-Rex1. A) N-terminal modules of P-Rex1 (gray, PDB: 8TUA, residues 38-499) and P-Rex2 (colored according to domain, residues 15-467) aligned to one another. There are minimal differences in their overall architecture. B) P-Rex2 aligned to P-Rex1 through the αH helix in the IP4P domain (P-Rex1 1491-1513, P-Rex2: 1431-1453) using Coot least-squares (LSQ) superimposition. A close-up view of each N-terminal domain (noted by the eyeballs) is shown in the insets, with P-Rex1 shown in gray. Spherical markers denote the center of mass (COM) of each homologous domain, and the distance between these in the aligned structures is indicated (PRex1 DH 38-237, PH 238-396, DEP1 397-499; P-Rex2 DH 15-211, PH 212-365, DEP1 366-467). C) The distance between the COM of the DEP1 domain and L1513/L1453 of the αH helix for P-Rex1 and P-Rex2, respectively, is shown. The measurement for an angle formed by a line connecting these two points and a line connecting L1513/L1453 to Q1491/A1431 of the αH helix for P-Rex1 and P-Rex2, respectively, is shown. Together, these describe the degree to which the N-terminal module is rotated and translated closer to the core in P-Rex2 as compared to P-Rex1.

Despite the overall structural similarity in P-Rex1 and P-Rex2, there is a significant shift in the N-terminal module relative to the core such that the P-Rex2 DEP1 domain is positioned proximal to the αH helix at the base of the core rather than distally to the side (Fig. 2B & C). To quantify this relative shift, the αH helices were aligned (RMSD Cα of 0.32 Å), and the centers of mass (COMs) of each N-terminal domain were used to measure the distances between homologous domain pairs in the N-terminus. Comparing P-Rex1 and P-Rex2, the DH, PH, and DEP1 domains are shifted by 19 Å, 30 Å, and 22 Å, respectively (Fig. 2B). The P-Rex2 DEP1 domain COM is ∼16 Å closer to the base of the αH helix than the P-Rex1 DEP1 COM (Fig. 2C). Additionally, the angle of the DEP1 COM from the αH helix decreases from 115° in P-Rex1 to 93° in P-Rex2, resulting in an overall more compact structure of P-Rex2 (Fig. 2C). Given this substantial relative shift in the N-terminal module, we mapped the existing P-Rex2 cross-linking mass spectrometry data^36^ onto the P-Rex2 structure to assess how well the data corroborated the N-terminal module position (Fig. S1). Distances between residues in the N-terminal module cross-linked to residues in the core are mostly within reason, based on the length of the cross-linker used, especially in comparison to the distances between homologous residues in P-Rex1 (Fig. S1). However, there are some exceptions. For example, K337 and K364 in the PH domain are too far away from K1442 in the IP4P domain. These cross-links also occur at low frequency in comparison to the others, which could suggest that these represent non-specific intermolecular interactions between different P-Rex2 molecules.

### Structural differences in the proposed phosphatase active site and Gβγ-binding elements

When the structure of the P-Rex1 C-terminal core was first determined in complex with Gβγ, it was discovered that the IP4P domain is structurally very similar to *Legionella* bacterial phosphoinositide 3-phosphatases^26^. Based on homology, the catalytic triad in P-Rex1 was proposed to be formed by Cys1583, Arg1589, and Asp1638 (Fig. S4E), although, to date, no substrates have been identified to confirm P-Rex1 phosphatase activity. Part of this “active site” is a loop containing Asp1638 that also forms a portion of the binding site for Gβγ, suggesting that any existing phosphatase activity could be dependent on Gβγ binding. P-Rex2 also shares this phosphatase fold; however, there are notable differences in the homologous residues that would form the catalytic triad. Although Cys1529 and Arg1535 are structurally conserved, Asp1584 is pointed away from the other triad residues, albeit map density in this area is weaker than in the rest of the “active site.” Furthermore, the loop on which Asp1584 resides is partially disordered and structurally different from the homologous loop in P-Rex1 where this loop is involved in Gβγ binding^26^. Intriguingly, Cys1529 seems to form a disulfide bond with Cys1536 even though the P-Rex2 cryo-EM sample was kept under reducing conditions at all times (Fig. S4F). The corresponding residue in P-Rex1 is Ser1590, which would not be able to form a disulfide bond, and this forms part of an a-helix that also appears to be involved in binding Gβγ^26^. Thus, there is potential for regulation of either phosphatase activity or Gβγ binding by disulfide bond formation. Because these two structural elements form major components of the Gβγ-binding site, these divergences could have implications for how P-Rex2 interacts differently with Gβγ as compared to P-Rex1.

### IP_4_ binding does not enable resolution of the P-Rex2 IP4P subdomain

In P-Rex1, the IP4P 4HB subdomain extends from the C-terminal core and makes contacts with the N-terminal module, promoting autoinhibition^24,25^. This closed, autoinhibited conformation is stabilized by binding of inositol-(1,3,4,5)-tetrakisphosphate (IP_4_), the soluble head group of the lipid PIP_3_, to the PH domain^25^. Addition of IP_4_ to the P-Rex1 cryo-EM sample facilitated resolution of both the 4HB and the N-terminal module by stabilizing the contact between the two, as these elements appear to be otherwise flexibly tethered to the core^26^. While the N-terminal module was clearly visualized in our P-Rex2 cryo-EM data, the IP4P subdomain was never resolved in any 2D class averages or 3D reconstructions despite extensive particle classification aimed at isolating subsets of particles where the subdomain might be observed (Fig. S2 & S3). The subdomain (∼29 kD) was confirmed to be intact in the P-Rex2 protein sample via molecular weight estimation by SDS-PAGE (Fig. S2A). Seeking to resolve the P-Rex2 IP4P subdomain, we added IP_4_ to the P-Rex2 cryo-EM sample and collected a small dataset. However, this did not alter the resulting 2D class averages or 3D reconstructions, and we were still unable to resolve the subdomain (Fig. S5). This suggests that IP_4_ binding does not affect the conformation of P-Rex2 as it does P-Rex1.

### Binding of IP_4_ has no effect on the tertiary structure of P-Rex2

To further examine the lack of effect of IP_4_ on visualization of the IP4P subdomain, we conducted hydrogen-deuterium exchange mass spectrometry (HDX-MS) on P-Rex2 in the presence and absence of IP_4_. HDX-MS measures deuterium uptake into a sample of protein on a per peptide basis, allowing assessment of the structure and dynamics of the protein. By comparing samples of P-Rex2 with and without IP_4_, we can assess how its structure changes upon IP_4_ binding. In P-Rex2, IP_4_ binding protects the β1/β2 and β6/β7 loops as well as the β6 strand from deuterium uptake through direct binding to the PIP_3_- binding site (Fig. 3A & B). Similarly, IP_4_ binding to P-Rex1 strongly decreases deuterium uptake in these structural elements (Fig. 3C; reanalyzed data from Ravala et al., 2024). However, it also decreases exchange in the P-Rex1 PH β5/β6 loop and IP4P 4HB_1_ and 4HB_2_ helices by stabilizing the interaction between the PH domain and the 4HB^25^ (Fig. 3C & E). Interestingly, the P-Rex2 PH β5/β6 loop also exhibits a strong decrease in deuterium uptake, yet P-Rex2 lacks any significant IP_4_-dependent protection within the IP4P subdomain or any other regions of the protein (Fig. 3A-B & D). This suggests either that IP_4_ has no effect on contacts between the N-terminal module and IP4P subdomain or that these interactions do not occur. Regardless, our HDX-MS data support that the P-Rex2 subdomain exhibits a structure very similar to the P-Rex1 4HB based on similarity of the deuterium uptake profiles in this region in the absence of IP_4_^25^ (Fig. S10).

**Figure 3.**
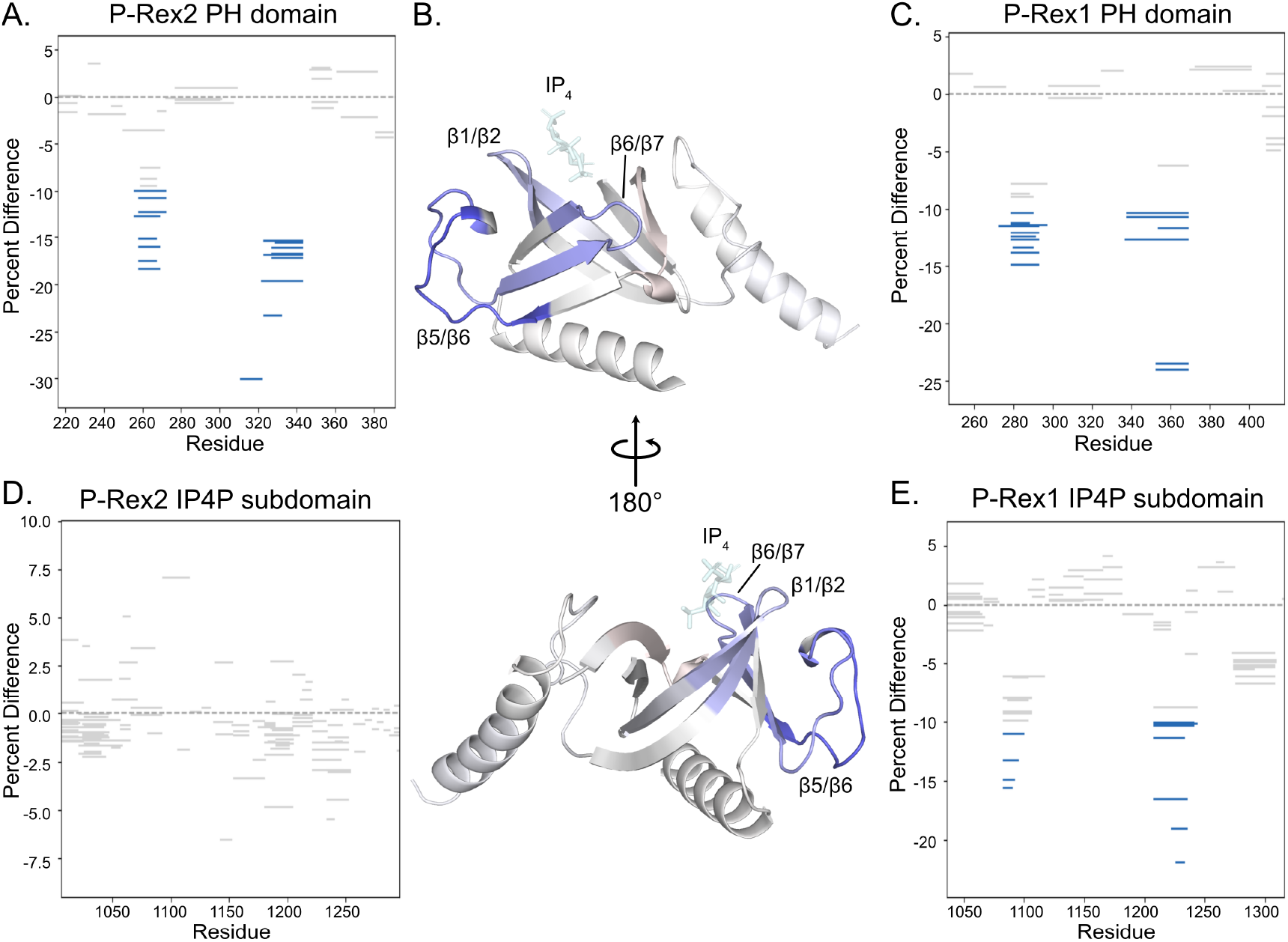
IP_4_ does not alter the dynamics of the P-Rex2 IP4P subdomain, unlike P-Rex1. HDX-MS data were collected on P-Rex2 in the presence and absence of IP_4_. A) HDX-MS peptide data at the 10,000 s time point shown on a Wood’s plot. Changes in protection greater than or equal to 10% are colored in blue. These changes indicate less deuterium uptake in the presence of IP_4_. The PH domain of P-Rex2 (216-381) exhibits IP_4_-dependent protection in the PIP_3_-binding site. B) P-Rex2 PH model colored according to difference HDX-MS data at the 10,000 s time point. Blue and light red regions indicate less and more deuterium uptake, respectively, in the presence of IP_4_. IP_4_ is docked based on the P-Rex1 PH•IP_4_structure (PDB: 5D3X) and shown transparently in cyan. Loops in between β-strands are labeled. C) HDX-MS data from Ravala et al., 2024 shown on a Wood’s plot for comparison. P-Rex1 PH (247-418) shows protection from deuterium uptake in the PIP_3_-binding site in the presence of IP_4_. D-E) The IP4P subdomain of P-Rex1 (1036-1316) shows increased protection from deuteration in the 4HB_1_ and 4HB_2_ helices. However, the P-Rex2 IP4P subdomain (996-1296) does not show any significant change in deuteration uptake in the presence of IP_4_.

### P-Rex1 is more active than P-Rex2 against Rac1

The distinct structural organization of P-Rex2 prompted us to investigate how its basal GEF activity might be different from that of P-Rex1. Utilizing a guanine-nucleotide exchange assay that measures GEF activity through loading soluble Rac1 with a fluorescently-tagged, nonhydrolyzable mant-GTP, we compared the activities of P-Rex2 and P-Rex1 constructs. We observed that a substantially higher concentration of FL P-Rex2 (300 nM) was required to achieve activity comparable to that of FL P-Rex1 (50 nM), supporting that P-Rex1 exchanges nucleotide on Rac1 more efficiently than P-Rex2 (Fig. 4A). To tease out whether higher GEF activity arises from differences in intrinsic catalytic efficiency or autoinhibition from domains outside of the catalytic core, we purified the isolated P-Rex DH/PH tandems and analyzed the activity of each (Fig. 4B & S6A). While P-Rex2 DH/PH displayed notably lower activity than the P-Rex1 DH/PH fragment, the difference in activities between the two was smaller, with the greatest difference in calculated k_obs_ at a single concentration being ∼3-4-fold as compared to the ∼6-fold higher concentration of FL P-Rex2 needed for equivalent activity. Together, these observations indicate that while the intrinsic catalytic activity of P-Rex2 DH/PH is weaker than that of P-Rex1 DH/PH, accessory domains in P-Rex2 FL also likely impose stronger autoinhibitory control of the catalytic tandem than in P-Rex1 FL. However, the mechanism of tighter regulatory control is unclear.

**Figure 4.**
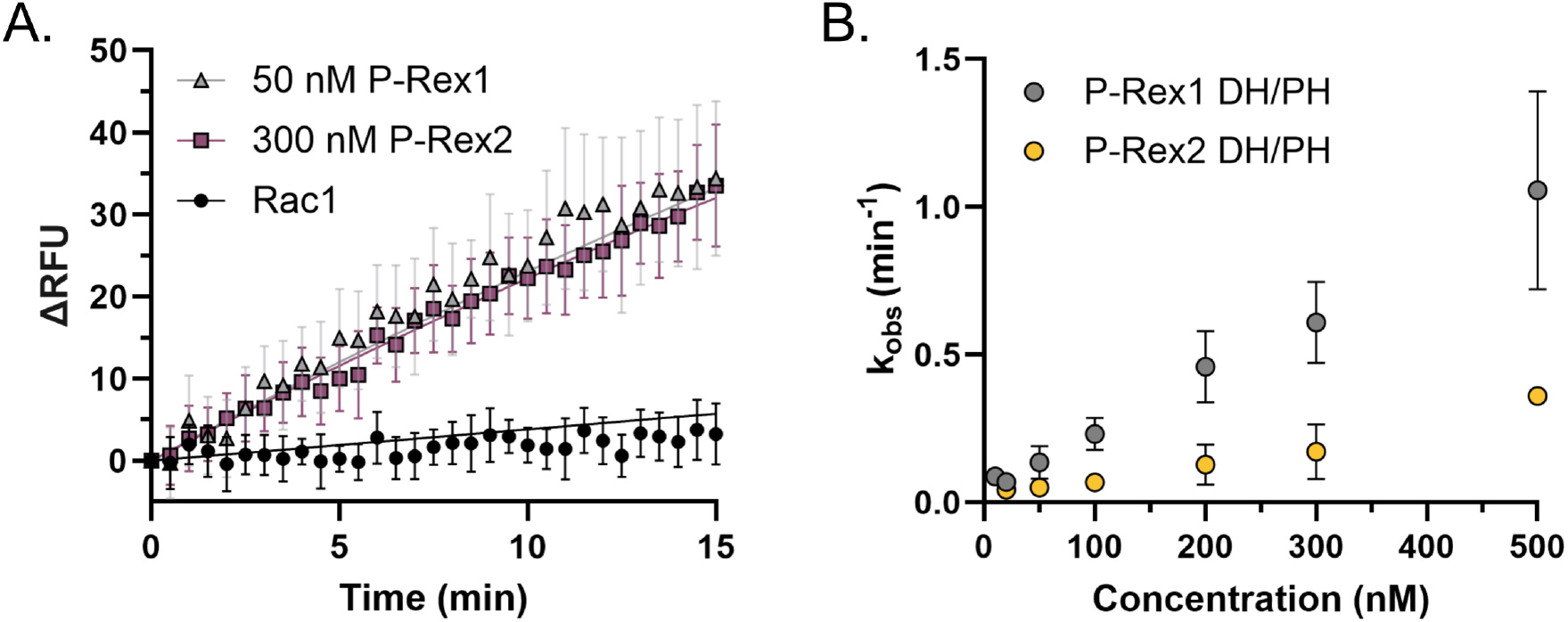
P-Rex2 exhibits lower basal activity against soluble Rac1 than P-Rex1. A) *In vitro* GEF activity assay with purified full-length P-Rex1 and P-Rex2 at 50 nM and 300 nM, respectively. B) GEF activity of purified P-Rex1 and P-Rex2 DH/PH constructs assessed at different concentrations. Error bars indicate standard deviation.

### The P-Rex2 DH/PH takes on a single, compact conformational state independent of DEP1

Previous size exclusion chromatography coupled with small-angle X-ray scattering (SEC-SAXS) experiments showed that the P-Rex1 DH/PH exists in two conformational states— the extended and compact form— whereas the DH/PH-DEP1 construct favors a single, compact conformation^27^. To determine if the P-Rex2 DEP1 domain also induces compaction of the catalytic core, we performed SEC-SAXS analysis on purified P-Rex2 DH/PH and DH/PH-DEP1 constructs (Fig. S6A). The SAXS profiles of both P-Rex2 constructs were consistent with monodisperse, well-folded particles (Fig. 5A & B, Table 2). Dimensionless Kratky analysis confirmed that both P-Rex2 DH/PH and DH/PH-DEP1 are compact and folded, exhibiting nearly identical profiles with well-defined peaks characteristic of globular proteins (Fig. 5C). Guinier analysis yielded correlation volumes corresponding to molecular weights of 41.6 and 57.1 kDa, respectively, within reasonable agreement with the expected molecular weights (44 and 51 kDa) (Fig. S6B & C). The pair distance distribution functions, P(r), indicated D_max_ values of 84 Å for P-Rex2 DH/PH and 105 Å for P-Rex2 DH/PH-DEP1 which suggests that the maximum particle dimension increases between the two constructs reasonably with the increase in mass (Fig. 5D). This is not the case in P-Rex1 where the maximum particle dimension of P-Rex1 DH/PH, ∼105 Å, is roughly the same as the DH/PH-DEP1, 104 Å^27^. Together, this suggests that the P-Rex2 DH/PH may not take on an extended state like the P-Rex1 DH/PH and that it favors compact states regardless of the presence of DEP1. To directly compare the data from P-Rex1 and P-Rex2, we analyzed the published P-Rex1 SAXS datasets (SASDHY9 and SASDHW9) and our P-Rex2 datasets using BilboMD^37^rather than the previously employed ensemble optimization method (EOM)^38^. Using the AlphaFold server to generate an initial model of each construct and BilboMD to produce ensemble models which represent highly populated conformations in solution, we identified that both P-Rex2 constructs were best fit by a single-state model (Fig. 5A & B). However, when we performed the BilboMD analysis on the P-Rex1 data (Fig. S7), BilboMD did not reproduce clear indication of two discrete conformations of the P-Rex1 DH/PH, as previously shown with EOM analysis^27^, and only suggested a more elongated state corresponding to a larger apparent molecular weight than would be expected for DH/PH. In all, these data suggest that both P-Rex2 DH/PH and DH/PH-DEP1 exist predominantly in compact, folded conformations in solution, indicating potentially reduced conformational flexibility as compared to P-Rex1.

**Table 2.** P-Rex2 SEC-SAXS data analyses.

|  | P-Rex2 DH/PH | P-Rex2 DH/PH-DEP1 |
| --- | --- | --- |
| **Guinier analysis** |  |  |
| R<sub>g</sub> (nm) | 26.12 +/- 0.38 | 29.11 +/- 0.47 |
| I(0) | 9.68 +/- 0.11 | 13.27 +/- 0.17 |
| Q<sub>min</sub> * R<sub>g</sub> | 0.455 | 0.684 |
| Q<sub>max</sub> * R<sub>g</sub> | 1.312 | 1.301 |
| **Molecular weight analysis** |  |  |
| Porod Volume (nm<sup>3</sup>) | 52300 | 77200 |
| MW (Vc) (kDa) | 41.6 | 57.1 |
| Shape (S&S) | Compact | Compact |
| D<sub>max</sub> (nm) | 81.1 | 96.0 |
| **P(r) analysis** |  |  |
| D<sub>max</sub> (nm) | 84 | 105.0 |
| R<sub>g</sub> (nm) | 26.94 +/- 0.16 | 29.99 +/- 0.28 |

**Figure 5.**
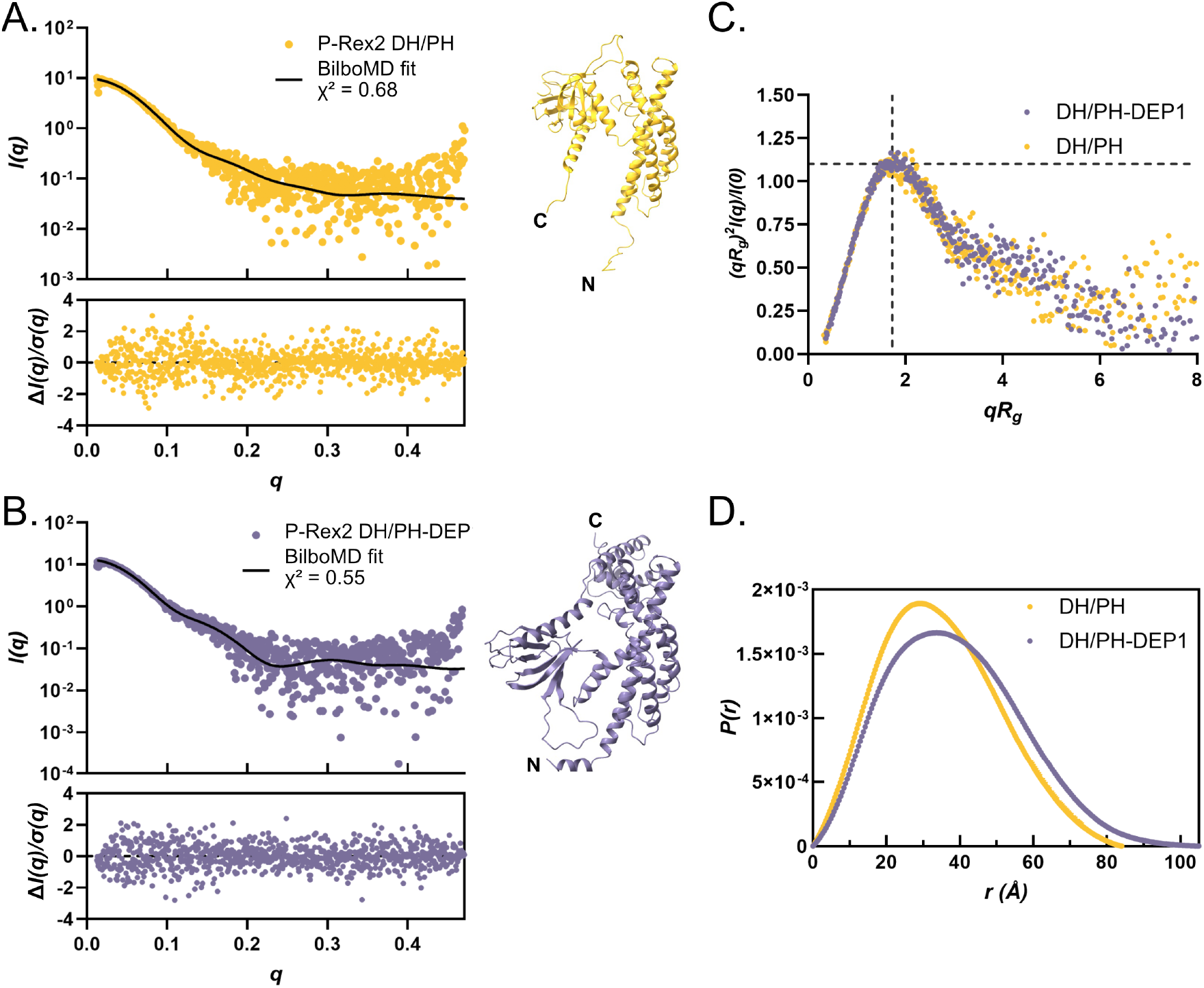
SEC-SAXS data show that P-Rex2 DH/PH takes on a single, compact state. A-B) Scattering intensity plots of P-Rex2 DH/PH (yellow) and DH/PH-DEP1 (purple) fit with the BilboMD model. The output BilboMD model is shown to the right of the plot. Normalized fit residuals are shown in the bottom panel. C) Dimensionless Kratky plot for P-Rex2 DH/PH and DH/PH-DEP1. There is no significant change in shape between the DH/PH and DH/PH-DEP1. D) Pair distance distribution function of P-Rex2 DH/PH and DH/PH-DEP1.

### The P-Rex2 DEP1 domain plays a role in maintaining autoinhibition

Previously, the DEP1 domain was shown to play an important role in maintaining P-Rex1 autoinhibition through contacts with the DH domain^24,25^. While P-Rex2 is structurally highly conserved within the N-terminal module, SEC-SAXS data alluded to potential differences in flexibility and conformation within the isolated DH/PH domain. This led us to investigate the role of DEP1 in P-Rex2 autoinhibition. We first compared the activities of P-Rex2 DH/PH, DH/PH-DEP1, and FL constructs using the guanine-nucleotide exchange assay with soluble Rac1 (Fig. 6A). As expected, P-Rex2 FL exhibited the lowest activity. The DH/PH-DEP1 construct exhibited intermediate activity between the full-length and DH/PH constructs, consistent with the DEP1 domain being involved in autoinhibition. Next, we introduced single point mutations in the DH/PH-DEP1 construct at the DH– DEP1 interface, focusing on homologous residues that are important for maintaining P-Rex1 autoinhibition^25^ and compared the activities of these variants to those of wild-type DH/PH-DEP1 and DH/PH (Fig. 6B-C & S8). Variants L151A, L152A, L434A, and L434F exhibited moderately increased activity, whereas I378A, I378F, and E424K showed the highest increases in activity. These data support that the hydrophobic pocket formed by Leu151, Leu152, Ile378, and Leu434 as well as a potential ionic contact at Glu424 collectively contribute to stabilization of the interface between the DH and DEP1 domains to promote autoinhibition (Fig. 6C). The K60A and K63A variants were generated due to the proximity of these residues to Glu424 in our predicted model before we obtained the cryo-EM structure of P-Rex2; however, we were unable to build these in our experimental model. The K60A variant did not express, and the L434 variants consistently exhibited low expression, which may reflect their importance in the stability of the N-terminal module (Fig. S8).

**Figure 6.**
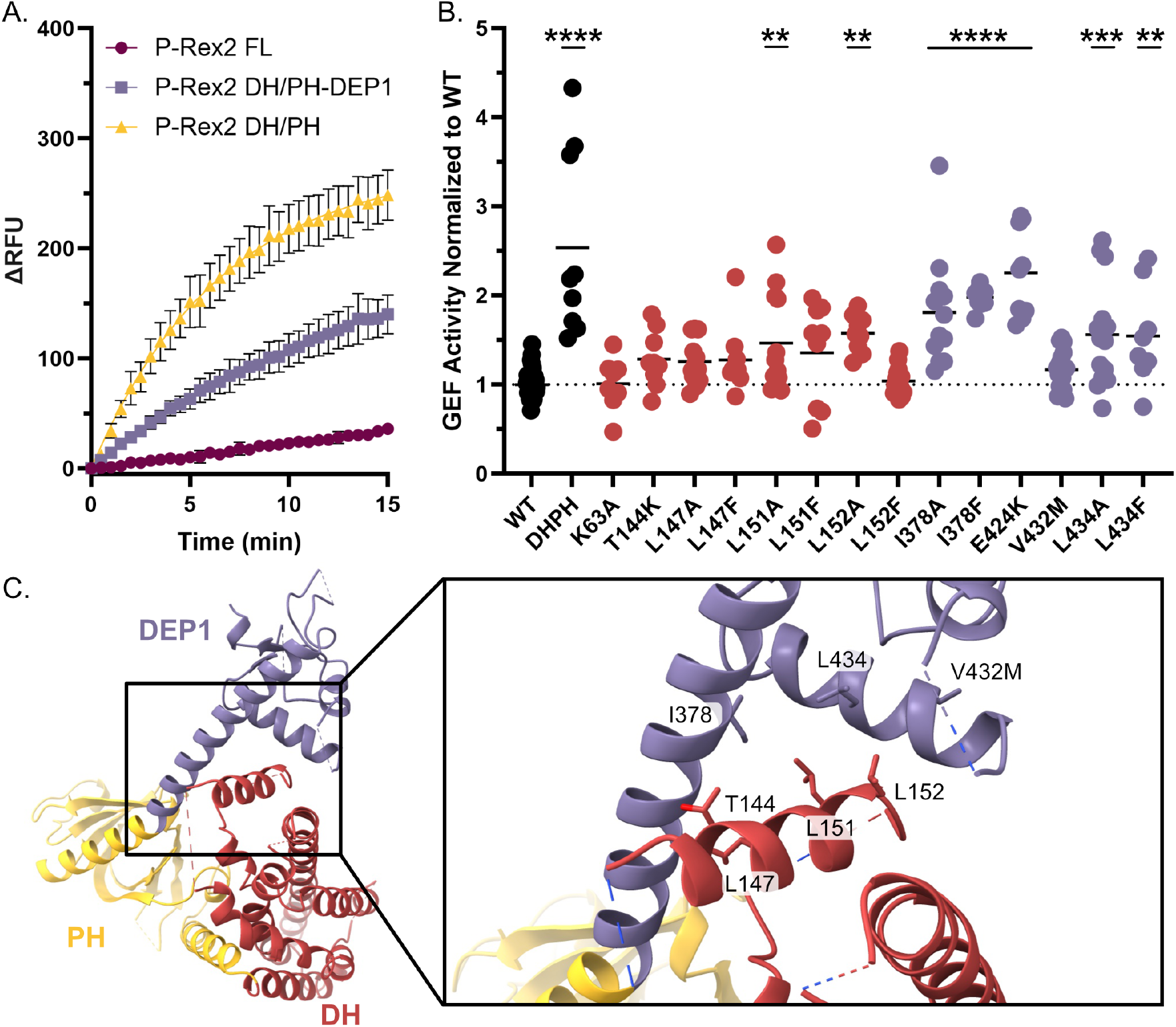
The P-Rex2 DEP1 domain exerts autoinhibition on the catalytic core via interactions with the DH domain. A) *In vitro* GEF activity assay with soluble Rac1 plus P-Rex2 FL (300 nM), DH/PH-DEP1 (300 nM), and DH/PH (200 nM). Error bars represent standard deviation. B) GEF activity assay with soluble Rac1 plus P-Rex2 DH/PH-DEP1 variants, colored according to the domain that the mutation lies in. C) Residues that were evaluated for their importance in maintaining autoinhibition through the DH–DEP1 interface.

## Discussion

Here, we presented the first structural characterization of autoinhibited P-Rex2 and revealed how its domain organization and regulatory features diverge from its close family member, P-Rex1 (Fig. 7). Our cryo-EM structure shows that while both the N- and C-termini are each individually structurally well-conserved between P-Rex1 and P-Rex2, the positions of these relative to one another are strikingly different between the two proteins. This major architectural dissimilarity may have significant consequences for how autoinhibition is achieved, pointing to potentially unique mechanisms being utilized in each isoform.

**Figure 7.**
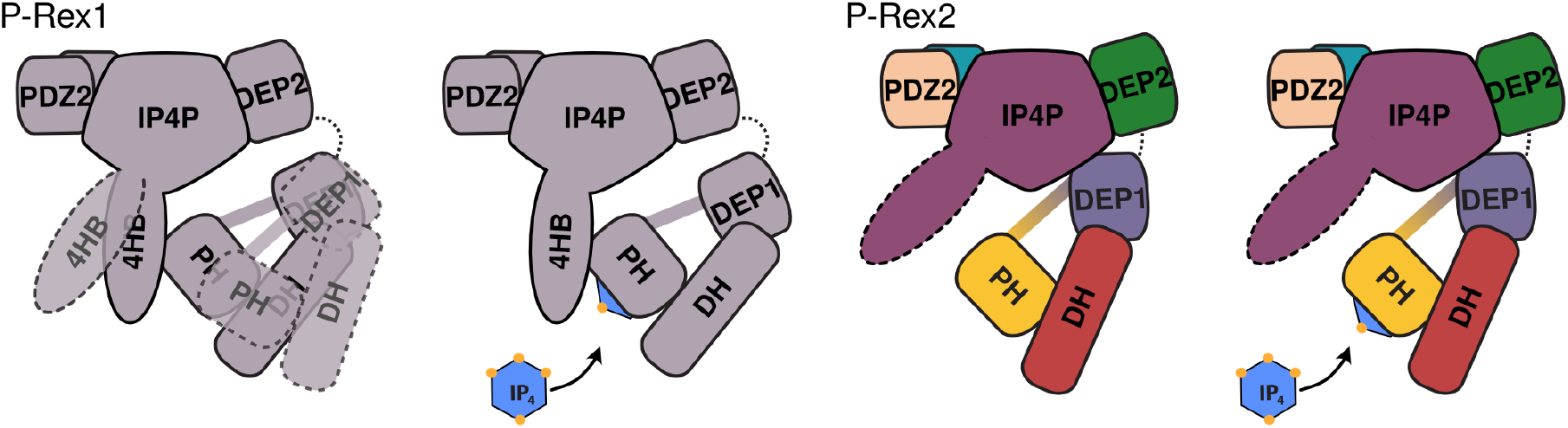
Overview of structural differences between P-Rex2 and P-Rex1. Although the structure of each the N-terminal module and C-terminal core are conserved between P-Rex2 and P-Rex1, the positioning of the two relative to one another is divergent. Furthermore, the autoinhibited form of P-Rex1 exhibits a conformation where the N-terminal module interacts with the 4HB subdomain of the IP4P domain, both of which appear to be otherwise flexibly tethered to the C-terminal core. This conformation is stabilized by binding IP_4_. In contrast, the P-Rex2 N-terminal module docks to the bottom of the C-terminal core and lacks any interaction with the IP4P subdomain which was not resolved in our cryo-EM data, presumably because it is flexibly tethered to the core. IP_4_ binding has no apparent effect on the structure or dynamics of P-Rex2 outside of the PIP_3_-binding site.

In P-Rex2, both the helix that spans the PH to DEP1 domains and the DEP1 domain itself are proximal to the base of the IP4P core. Docking of the N-terminal module to the core likely enables its resolution in our cryo-EM structure by tethering the otherwise likely flexible N-terminus. However, it is difficult to pinpoint the underlying cause of this structural difference between P-Rex proteins. The surface of the P-Rex2 N-terminal module that interacts with the core is very similar to that of P-Rex1, suggesting that this is not the source of the divergence. One hypothesis is that the connection between the first and second DEP domains in P-Rex2 is more rigid and inflexible than that in P-Rex1, as this linker is composed of bulkier, less flexible residues in P-Rex2 (TFYPRNE) compared to P-Rex1 (TYKARSE). This is supported by clear map density representing the P-Rex2 DEP1-DEP2 connection, although it is not high enough resolution for us to be able to confidently model into it. Alternatively, the unique structural features at the base of the P-Rex2 core could promote the docking of the N-terminal module. In P-Rex1, the IP4P αH and αI helices are moderately well-defined and are connected by a disordered loop of ∼14 residues (Fig. S9A). In P-Rex2, the homologous residues are much more well-resolved, forming a helical structure that packs very closely to the αH helix (Fig. S9B & C). Although the functional consequences of these structural features remain unclear, elucidating whether these elements contribute to autoinhibition could provide insight into regulatory surfaces that could potentially be exploited for P-Rex-specific inhibition.

Contrary to the P-Rex1 autoinhibitory mechanism, P-Rex2 does not exhibit a PH–IP4P subdomain interaction, with or without IP_4_. While our HDX-MS data suggest that the P-Rex2 IP4P subdomain may take on a similar tertiary structure as P-Rex1, the position of the N-terminal module makes it unlikely that the IP4P subdomain would be able to make the same contacts with the PH domain, as the P-Rex2 PH domain is positioned farther away from the C-terminal core. Although our present data do not support a clear role for the P-Rex2 IP4P domain in autoinhibition, it may help contextualize data pointing to a role in PTEN-induced inhibition. Cross-linking mass spectrometry of P-Rex2^36^ shows limited intramolecular contacts between the IP4P subdomain and the N-terminal module. However, upon PTEN binding, there is a substantial increase in the number of cross links between the IP4P subdomain and the DH and PH domains. This raises the possibility that PTEN does not primarily inhibit P-Rex2 through direct engagement with the N-terminus as previously suggested^10^, but instead binds to the PDZ2 and IP4P subdomain^36^ to strengthen long-range interdomain interactions between the IP4P and DH/PH. In this context, it is notable that the pancreatic cancer-associated V432M mutation, which lies near the DH–DEP1 interface, escapes PTEN-driven inhibition in breast cancer cells despite retaining PTEN binding^39^. Our biochemical assays using the isolated N-terminal module show only a minimal, non-significant increase in basal nucleotide exchange activity in the V432M variant, suggesting that this mutation does not drive disease through disruptions to autoinhibition from DEP1. However, these data imply that, in the full-length protein, the V432M mutation potentially changes how the N-terminal module interacts with the IP4P subdomain to disrupt PTEN-driven inhibition. While our understanding of the P-Rex2–PTEN coinhibitory complex is still limited, it would be valuable to investigate this mechanism further to identify how disruptions to this complex contribute to cancers and diabetes and, furthermore, to tease out why PTEN is a regulator of P-Rex2 and not P-Rex1.

We observed that the isolated P-Rex2 DH/PH exhibits slower nucleotide exchange on soluble Rac1 than that of P-Rex1. We also showed that, of the two identified mechanisms that govern P-Rex1 autoinhibition, only the DH–DEP1 interface is conserved in P-Rex2. However, the DH–DEP1 interface alone does not fully account for autoinhibition in P-Rex2, as P-Rex2 FL exhibits significantly lower activity than DH/PH-DEP1. This suggests that other regions of the protein must also participate in achieving full autoinhibition. Further examination of the C-terminal accessory domains for their possible roles in autoinhibition is needed. Additionally, biochemical and biological exploration of the alternative spliceoform P-Rex2b, which largely lacks the IP4P domain, has been very limited but might provide insight into a P-Rex2 that more closely exhibits the higher activity of P-Rex1.

Because of the unexpected differences in structure and autoinhibitory regulatory mechanisms, it is possible that P-Rex1 and P-Rex2 respond differently to activation by PIP_3_ and Gβγ. For example, interactions between the P-Rex1 PH domain and IP4P subdomain contribute to autoinhibition and must be disrupted for activation^25^, whereas the lack of such coupling in P-Rex2 implies that the PH domain may be more readily available to bind to PIP_3_, allowing P-Rex2 to become activated at lower PIP_3_ concentrations despite exhibiting lower basal activity than P-Rex1. There are also potential differences in Gβγ binding. In P-Rex1, the region around Gln1615-Leu1647 was identified to play a major role in binding Gβγ^26^. Within the homologous area in P-Rex2 (Lys1578-Pro1587), the structure diverges significantly. In another P-Rex1 element implicated in Gβγ binding (Met1582-Val1594), the homologous region in P-Rex2 appears to form an unusual disulfide bond that cannot form in P-Rex1 (Fig. S4E & F). These differences suggest that P-Rex2 may bind Gβγ with a different mechanism and/or affinity than P-Rex1, potentially eliciting a distinct activation response. However, our understanding of the activation mechanisms of P-Rex RhoGEFs is still largely unexplored. Future work investigating this is still needed to understand how P-Rex1 and P-Rex2 escape their individual autoinhibitory mechanisms.

In summary, this work has begun to uncover how P-Rex2 regulation is unique from P-Rex1 and how its structural elements could contribute to these differences. Although both proteins share high sequence similarity, P-Rex1 and P-Rex2 autoinhibition are achieved in part through distinct domain organization. This highlights opportunities to selectively target one isoform over the other through divergent regulatory surfaces outside of the DH domain via the rational design of modulators. Attempts at specifically targeting individual Dbl RhoGEFs have historically been unsuccessful due to the flat, broad DH domain that is highly conserved and binds structurally similar GTPases. More broadly, these findings are exciting because they are proof of principle that even closely related Dbl RhoGEFs within the same subfamily can exhibit clear differences in their structure and regulation, offering a framework for designing selective modulators that target Dbl RhoGEFs by utilizing their accessory domains. Future efforts integrating structural and cellular studies will be essential for translating these mechanistic insights into strategies for therapeutically targeting misregulated RhoGEF signaling and, furthermore, fine-tuning GTPase signaling to treat and prevent disease.

### Experimental procedures

#### Sequence Identity Comparison to P-Rex1

Using our P-Rex2 model, we identified the residue boundaries of each of the seven domains: DH: 15-211, PH: 212-365, DEP1: 366-467, DEP2: 473-569, PDZ1: 590-672, PDZ2: 673-815, and IP4P: 816-1606. The corresponding domain boundaries were identified in P-Rex1 based on structure comparison. Sequence identity was calculated using The Sequence Manipulation Suite^40^.

#### Cloning and plasmids

P-Rex1 full-length (FL), P-Rex1 DH/PH, and Rac1 DNA constructs were described previously^5,26,27^. P-Rex2 pcDNA3.1/V5-His DNA was a gift from Ramon Parsons (Addgene plasmid #41555)^35^. P-Rex2 FL was cloned into a pRK5 mammalian expression vector with a TEV-cleavable N-terminal GST tag and a C-terminal FLAG tag. P-Rex2 DH/PH (residues 1-377) and P-Rex2 DH/PH-DEP1 (residues 1-467) were each cloned into the pMALc2H_10_T vector with a TEV-cleavable N-terminal MBP-10xHis tag. P-Rex2 DH/PH-DEP1 point mutations were generated using site-directed mutagenesis.

#### Protein Production and Purification

P-Rex1 and P-Rex2 FL were expressed in FreeStyle 293-F cells by transient transfection using polyethylenimine. Cells were harvested 48 hours post-transfection by centrifugation for 15 min at 1,000 x g, flash frozen in liquid nitrogen, and stored at −80 °C. Pellets were thawed and lysed in Cell Lytic M (Sigma) supplemented with 200 mM NaCl, 1 mM ethylenediaminetetraacetic acid (EDTA), 2 mM dithiothreitol (DTT), and a protease inhibitor cocktail (125 nM aprotinin, 6.8 µM leupeptin, 0.15 nM soybean trypsin inhibitor, 1 µM E-64, and 1 µM bestatin). After rocking at 4 °C for 15 min, lysates were clarified by ultracentrifugation at 40,000 rpm for 45 min at 4 °C in a Type 45 or 70 Ti rotor depending on the size of the sample. The supernatant was filtered through a glass fiber filter and incubated with 800 µL glutathione agarose resin per liter harvested cells for 1-2 hours at 4 °C with gentle rocking. The resin was washed with 20 mM HEPES (pH 8), 200 mM NaCl, 1 mM EDTA, 2 mM

DTT, and protease inhibitor cocktail. GST-tagged P-Rex proteins were eluted with 100 mM HEPES (pH 8), 300 mM NaCl, 1 mM EDTA, 1 mM DTT and 30 mM reduced glutathione. Overnight at 4°C, elution fractions were digested with TEV protease (1:1 molar ratio) and dialyzed against 20 mM HEPES (pH 8), 100 mM NaCl, 1 mM EDTA, and 2 mM DTT. Protein was then further purified over an ENrichQ (BioRad) anion exchange column in 20 mM HEPES (pH 8), 2 mM DTT and eluted with a NaCl gradient from 100 mM to 550 mM over 30 CV. Purified protein was pooled, concentrated, and immediately used for experiments.

Soluble Rac1 was expressed in Rosetta (DE3) pLysS *E. coli* cells and purified as previously described^5^. Truncated P-Rex constructs also were expressed in Rosetta (DE3) pLysS *E. coli* cells as N-terminal His-tagged MBP fusion proteins. Cells were grown in Terrific Broth to an OD_600_ of 0.8, induced with 0.1 mM IPTG at 16 °C, harvested after 16-20 hours, and then flash frozen and stored at −80 °C. Cell pellets were thawed and resuspended in 9 ml 20 mM HEPES (pH 8), 300 mM NaCl, 0.1 mM EDTA, 2 mM DTT and protease inhibitor cocktail per 1 g cell pellet. Cells were then homogenized with a dounce and lysed with an Avestin EmulsiFlex-C5 High Pressure Homogenizer. Lysate was clarified by ultracentrifugation at 40,000 rpm for 45 min in a Type 45 Ti rotor. The supernatants were then filtered through a glass fiber filter and incubated with Ni-NTA resin for 30-60 min. Resin was washed with buffer (20 mM HEPES pH 8, 300 mM NaCl, 2 mM DTT) followed by buffer containing 10 mM imidazole, and proteins were eluted with buffer containing 250 mM imidazole. DH/PH and DH/PH-DEP1 elutions were then simultaneously dialyzed into buffer containing 20 mM HEPES (pH 7 or pH 8, respectively), 200 mM NaCl, 2 mM DTT, and 10% glycerol and digested with TEV protease overnight at a 1:2 molar ratio of TEV:MBP-fusion protein to remove the N-terminal His-tagged MBP. Cleaved MBP-His was then captured by an additional pass over Ni-NTA resin. The flow through was applied to a HiTrap SP Sepharose Fast Flow column (Cytiva) in 20 mM HEPES (pH 7) and 2 mM DTT and eluted over an NaCl gradient from 0-0.5 M over 4 CV.

P-Rex2 DH/PH-DEP1 mutants were expressed under the same conditions as the wild-type construct, but in small-scale 15 ml cultures. Cell pellets were resuspended in 1 ml 20 mM HEPES (pH 8), 300 mM NaCl, 0.1 mM EDTA, 2 mM DTT, and protease inhibitor cocktail. Resuspended cells were sonicated on ice (5 seconds on, 5 seconds off, for 60 seconds per cycle for 3 cycles) using a Q125 sonicator (Qsonica Sonicators), and lysates were clarified by centrifugation at 21,300 x g for 45 min at 4°C. The supernatants were incubated with Ni-NTA resin for 30-60 min at 4°C. The resin was washed and proteins eluted, cut with TEV, and dialyzed the same as the wild-type DH/PH-DEP1. Cleaved MBP-His was then captured by an additional pass over Ni-NTA resin. The resin was washed one additional time with a buffer containing 20 mM HEPES (pH 8), 300 mM NaCl, 2 mM DTT, and 10 mM imidazole. The combined flow through and wash were concentrated using a 10kD MWCO Amicon concentrator. Final protein concentrations were estimated by visualization on SDS-PAGE.

#### Cryo-EM Grid Preparation and Data Acquisition

P-Rex2 samples at 0.4 mg/ml were briefly incubated with 0.07 mM n-Dodecyl-Beta-Maltoside (DDM) before applying a 4 µL sample to a glow discharged 300-mesh Quantifoil (1.2/1.3) grid, blotted for 4 or 6 seconds with a blot force of 20, and plunge-frozen into ethane cooled with liquid nitrogen using a Vitrobot (Thermo Fisher Scientific) set to 4 °C and 100% humidity. Data were collected at the Pacific Northwest Center for Cryo-EM (PNCC) using EPU (Thermo Fisher) on a Krios transmission electron microscope operating at 300 keV with a Falcon 4i direct electron detector (Thermo Fisher) and SelectrisX. Datasets were collected on an untilted grid (12,070 micrographs) with beam image shift (BIS) and on a grid tilted by 35° (7,722 micrographs) without BIS. Tilted data were collected to overcome a preferred orientation of the sample on grids.

For the IP_4_ experiments, small datasets were collected on P-Rex2 samples at 0.7 mg/ml that were briefly incubated with 0.07 mM DDM in the presence or absence of 16.6 µM IP_4_ and frozen with the same freezing conditions described above. Approximately 1300-1400 micrographs were collected for each dataset using a Glacios transmission electron microscope operating at 200 keV with a K3 direct electron detector (Gatan, Inc.) and then processed up through 2D classification. Both samples yielded classes with similar views without distinguishable differences in P-Rex2 conformation, mass present, or resolution (Fig. S5).

#### Cryo-EM Data Processing

Untilted and tilted datasets were preprocessed separately. Motion correction, contrast transfer function (CTF) estimation, and particle picking were performed in Warp^41^. The default BoxNet2Mask model was re-trained on P-Rex2 particles for particle picking in these datasets. Particle stacks were then imported into cryoSPARC^42^ for further processing. Each particle stack was cleaned through multiple rounds of 2D classification and then stacks from the untitled and tilted data were combined and used for ab-initio reconstruction (Fig. S3).

In the main processing pipeline (Fig. S3), ab-initio reconstruction yielded one volume where both the C-terminal core and the N-terminal module were resolved and 3 volumes where mainly only the core was resolved. Only the volume with clear density for the N-terminal module was used for downstream processing. This volume was processed through non-uniform refinement^43^, yielding a 3.4 Å reconstruction according to cryoSPARC. This reconstruction along with three decoy volumes created from “junk” particles were used in heterogeneous refinement against the cleaned particle stack, resulting in ∼100k additional particles sorting into the “good” class. Additional non-uniform refinement of this stack and volume resulted in a 2.9 Å consensus map which featured primarily only the core (Fig. S2D). To resolve the N-terminal module, focused 3D classification was performed on the consensus map and particles using a mask around the N-terminal module, and one class from this was then locally refined using the same mask (Fig. S2H). The consensus map and the map from local refinement of the N-terminal module were used to generate a composite map in Phenix, utilizing a 4.5 Å resolution cutoff (Fig. S4D).

In a completely alternative processing pipeline (not shown), starting with a cleaned particle stack, several “good” and decoy volumes were generated by ab-initio reconstruction and used to begin multiple rounds of heterogeneous refinement to sort out particles that would produce reconstructions containing both the core and N-terminal module, followed by non-uniform refinement. Then, the best volume and particles from this were put through 3D refinement with Blush regularization in Relion 5.0^44^. Finally, the refined map and particles underwent local refinement in cryoSPARC with a mask around the whole particle. The resulting map was put through density modification using EMReady^45^ to improve interpretability of the map for use in model building (Fig. S4B & C).

#### Model Building, Refinement, and Validation

Primarily, two maps were used to build the P-Rex2 model, as each map was more interpretable in different areas. One of these was a composite map and the other was a density-modified map from EMReady, both of which are described above. Using the AlphaFold Server^46^, P-Rex2 models of each the N-terminus (residues 1-472) and C-terminus (473-1606) were generated. Models were fit into the aligned maps using ChimeraX^47^, and the best fitting model for each the N- and C-terminus was chosen for further model building and modification. Parts of the model that were completely outside of map density were deleted in Coot 0.9.8.95^48^, and then each model was run through rigid body real-space refinement in Phenix 1.21.2-5419^49^ after adding hydrogens and with minimization_global enabled^50^. After this, the model was further modified and rebuilt in Coot using tight geometric restraints. Because map density was very sparse or low resolution in some regions, homology models P-Rex1–Gβγ (6PCV), P-Rex1•IP4 (8TUA), and P-Rex1 DH/PH– DEP1 (7RX9) were used to assist decision making during model building. In some areas, the AlphaFold model was different from the homologous region in P-Rex1 but nicely fit the P-Rex2 map, while in others, the predicted and homology models clearly deviated from the path of the map. In regions where the map was ambiguous, the AlphaFold model was left with limited modification when it fit the density reasonably well, corresponded to helical or β-strand regions, and was conserved in P-Rex1. In areas where the P-Rex2 map diverged from P-Rex1, such as in loops where the prediction was less reliable or the density path was not clear, the model was deleted. After building was complete, regions of the structure that formed helices or β-sheets were manually confirmed and annotated, and Phenix real-space refinement was run using secondary structure restraints. Model validation was performed in Phenix^50^ (Table 1 & Fig. S4A). EMRinger was also run within Phenix^51^. The associated web interface was used to run the 3D FSC program to calculate sphericity and generate a 3D FSC histogram and directional FSC plot^52,53^. The Q score was calculated during the validation stage of data deposition^54^. Structure figures were generated using ChimeraX^47^ and PyMOL^55^. Some software was accessed through membership in the SBGrid Consortium^56^.

#### Hydrogen-Deuterium Exchange Mass Spectrometry

Prior to conducting hydrogen/deuterium exchange experiments, optimal quench conditions were determined to generate the best enzymatic peptide coverage map for P-Rex2 FL, as previously described^26,57^. Briefly, 3 μl purified P-Rex2 (2 mg/ml in 20 mM HEPES (pH 8), 150 mM NaCl, 2 mM DTT) was diluted with 9 μl of H_2_O Buffer at 0 °C, and then mixed with 18 μl of ice cold quench buffers containing 0.1 M Glycine (pH 2.4), 16.6% glycerol and varying concentration of GuHCl (0.08, 0.8, 1.6 and 3.2 M). The quenched samples were subjected to on-line proteolysis and LC/MS analysis. The best sequence coverage of P-Rex2 was obtained using the 1.6 M GuHCl quench buffer.

P-Rex2 exchange stock solutions (10.8 μM of P-Rex2 FL and 10.8 μM of P-Rex2 FL with 45 μM IP_4_) were prepared in 8.3 mM Tris, 150 mM NaCl, pH 7.2 and kept on ice. HDX experiments were initiated by adding 50 μl of exchange stock solutions (P-Rex2 FL or P-Rex2 FL-IP_4_) to 150 μl of D_2_O buffer (8.3 mM Tris, 150 mM NaCl, pDread7.2) and incubating for various times intervals (10, 100, 1000, 10,000 and 100,000 sec) at 0 °C. At each time, 36 μl of exchange mixture was withdrawn and quenched with 54 μl of ice cold quench buffer (0.1 M Glycine (pH 2.4), 1.6 M GuHCl). Quenched samples were aliquoted and immediately frozen on dry ice. Non-deuterated and equilibrium deuterated control samples were also prepared for back exchange correction^58^.

All frozen samples were later loaded onto a cryogenic autosampler and automatically thawed at 4 °C before digestion on an immobilized pepsin column (16 μl bed volume) at 25 μl/min. Proteolytic fragments were collected on a trap column and separated on an Acclaim PepMap RSLC C18 reverse phase analytical column (ThermoSci, 0.3 x 50 mm, 2 μm, 100 Å) using a linear acetonitrile gradient (5%-45% over 30 min).

The effluent was directed into an OrbiTrap Elite Mass Spectrometer (ThermoFisher Scientific, San Jose, CA). Instruments settings were optimized to minimize the back-exchange^59^. The data was acquired in either MS1 profile mode or data-dependent MS/MS mode. Peptide identification was done with the aid of Proteome Discoverer software (ThermoFisher). The centroids of the mass envelopes of deuterated peptides were calculated with HDExaminer (Sierra Analytics Inc, Modesto, CA) and then converted to corresponding deuterium incorporation with corrections for back-exchange^60^.

#### GTPase Exchange Activation Assay

P-Rex GEF activity was analyzed through a FRET-based nucleotide exchange assay measuring association of GTPase with fluorescently-labeled N-methyl-anthraniloyl-GTP (mant-GTP; Abcam), a nonhydrolyzable GTP analog, in a black 384-well plate (Greiner). P-Rex construct plus 2 µM GTPase were added in 20 mM HEPES (pH 8), 100 mM NaCl, 5 mM MgCl_2_, 1 mM DTT, and 5% glycerol at room temperature. The reaction was initiated by adding 0.8 mM mant-GTP and immediately measured for fluorescence at Ex = 280 nm, Em = 450 nm at room temperature in 30 second intervals for 15 min on a SpectraMax M5 (Molecular Devices) plate reader. Fluorescence curves were then fitted to a one-phase association model using GraphPad Prism.

#### Statistical Analysis

All statistical analyses were performed in GraphPad Prism software on data from at least three independent experiments. All GEF activity assay data were fit to a one-phase association model. The k value obtained was used to represent k_obs_ where applicable. All P-Rex2 DH/PH-DEP1 variants were normalized to P-Rex2 DH/PH-DEP1 WT and then evaluated for statistical significance. Statistical significance was determined using a one-way ANOVA test with a post hoc Dunnett’s test for multiple comparisons. * p < 0.05, ** p < 0.01, **** p < 0.0001.

#### Size-Exclusion Chromatography Small-Angle X-ray Scattering

Small-angle X-ray scattering (SAXS) experiments were performed at the SIBYLS beamline 12.3.1 at the Advanced Light Source. This beamline has additional inline instrumentation and detectors coupled to a size exclusion column^61,62^. Purified P-Rex2 DH/PH and DH/PH-DEP1 were concentrated to 2 mg/ml and then buffer exchanged into 20mM HEPES (pH 7), 200mM NaCl, 2mM DTT, and 1% glycerol SEC running buffer. The X-ray wavelength was set to 1.127 Å and the sample-to-detector distance to 2,100 mm, which gives scattering vectors (q) ranging from 0.01 Å^-1^ to 0.4 Å^-1^. The scattering vector is q = 4πsinθ/λ, where 2θ is the scattering angle. The SAXS flow cell was coupled to an Agilent 1260 Infinity HPLC system using a Shodex PROTEIN KW-803 SEC column equilibrated with the running buffer with a flow rate of 0.65 ml/min. For SAXS measurements, 2 sec X-ray exposures were collected continuously during a 25 min elution. All frames for analyses had one SAXS frame corresponding to the running buffer before the detection of a peak subtracted from each.

The radius of gyration (Rg) was calculated for each of the subtracted frames using the Guinier approximation: I(q) = I(0) exp(−q2Rg2/3) with the limits qRg < 1.3. The elution peak was compared to the integral of ratios to background and Rg relative to the recorded frame using the program RAW^63^. Uniform Rg values across an elution peak represent a homogeneous sample. Final merged SAXS profiles, derived by integrating multiple frames at the elution peak, were used for further analyses. Calculated were the Guinier plot to provide information on the aggregation state, the volume of correlation (Vc) to estimate the molecular weight^64^, and the pair distribution function [P(r)] to calculate the maximal inter-particle dimension^65^. AlphaFold Server^46^ was used to generate models of P-Rex1 DH/PH (38-409), P-Rex2 DH/PH (1-377), P-Rex1 DH/PH-DEP1 (38-499) and P-Rex2 DH/PH-DEP1 (1-467). The resulting models were used with their corresponding intensity profiles in BilboMD^37^ to determine the model fit to the intensity profile and potential conformational heterogeneity.

## Data availability

The P-Rex2 EM data can be accessed as EMDB entries EMD-74547 (consensus map), EMD-74548 (local refinement of the N-terminal module), EMD-74549 (local refinement of the whole particle), EMD-74550 (composite map) and EMPIAR entry EMPIAR-XXXX. The P-Rex2 model has been deposited as PDB entry 9ZQ7. The SAXS data for P-Rex2 DH/PH and DH/PH-DEP1 can be accessed as SASDB entry XXXX and XXXX, respectively.

## Supporting information

This article contains supporting information.

## Acknowledgements

We would like to thank Joshua Del Mundo and Michal Hammel at the Lawrence Berkeley National Laboratory for their assistance with SEC-SAXS data interpretation. We would also like to thank Fei Guo for support in the UC Davis BioEM facility and Hemang Patel and Camille Scott for IT and HPC support for cryo-EM data storage and processing. We furthermore would like to thank Emma Barpal for providing technical support for the project.

## Author contributions

C.P. expressed FL P-Rex proteins and banked cell pellets, assisted with purification of these proteins, and performed initial assays with these proteins. R.M. cloned the P-Rex2 DH/PH and DH/PH-DEP1 constructs and performed initial test expressions. V.N. and G.M. cloned P-Rex2 DH/PH-DEP1 variants, expressed these variants, and assisted with some of their purifications. S.L. performed HDX-MS experiments on purified P-Rex2 samples and analysis of the raw data. L.K.A. performed all other benchwork, including preparing samples for HDX-MS and SEC-SAXS experiments and preparing and screening samples for cryo-EM, processed cryo-EM data, analyzed experimental results, prepared figures, and wrote the initial draft of the manuscript.

## CRediT author contributions

Conceptualization - L.K.A and J.N.C.

Data curation - L.K.A, J.N.C., S.L.

Formal analysis - L.K.A, J.N.C., S.L.

Funding acquisition - J.N.C.

Investigation - all authors

Methodology - L.K.A and J.N.C.

Project administration - J.N.C.

Resources - J.N.C.

Supervision - L.K.A and J.N.C.

Validation - L.K.A and J.N.C.

Visualization - L.K.A and J.N.C.

Writing — original draft - L.K.A and J.N.C.

Writing — review & editing - all authors

## Funding and additional information

Research reported in this publication was supported by the National Institute of General Medical Sciences of the National Institutes of Health under Award Number R35GM146664 (to J.N.C.). This project was supported in part by Administrative Coordinating Council of Deans (ACCD) College of Biological Sciences funds received from the UC Davis Clinical and Translational Science Center, which is supported by award UL1 TR001860 from the NIH National Center for Advancing Translational Sciences (to J.N.C.). A portion of this research was supported by NIH grant R24GM154185 and performed at the Pacific Northwest Center for Cryo-EM (PNCC) with assistance from Marcelo De Farias. SAXS data was collected at the Advanced Light Source (ALS) at the SIBYLS beamline, a national user facility operated by Lawrence Berkeley National Laboratory on behalf of the Department of Energy, Office of Basic Energy Sciences, through the Integrated Diffraction Analysis Technologies (IDAT) program, supported by DOE Office of Biological and Environmental Research. Additional support comes from the National Institute of Health project ALS-ENABLE (P30 GM124169). Molecular graphics and analyses performed with UCSF ChimeraX, developed by the Resource for Biocomputing, Visualization, and Informatics at the University of California, San Francisco, with support from National Institutes of Health R01-GM129325 and the Office of Cyber Infrastructure and Computational Biology, National Institute of Allergy and Infectious Diseases. The content is solely the responsibility of the authors and does not necessarily represent the official views of the National Institutes of Health.

## Abbreviations and nomenclature

RhoGEF: Rho guanine-nucleotide exchange factor
PIP_3_: phosphatidylinositol-3,4,5-trisphosphate
P-Rex: phosphatidylinositol-3,4,5-trisphosphate-dependent Rac exchanger
GDP: guanosine diphosphate
GTP: guanosine triphosphate
Rac: Ras-related C3 botulinum toxin
Cdc42: cell division control protein 42 homolog
Dbl: diffuse B-cell lymphoma
DH: Dbl homology
PH: pleckstrin homology
DEP: Dishevelled, Egl-10 and Pleckstrin
PDZ: Postsynaptic density protein 95, Discs large protein, and Zonula occludens-1
IP4P: inositol polyphosphate-4-phosphatase-like
PI3K: phosphoinositide 3-kinases
4HB: 4-helix bundle
IP_4_: inositol-(1,3,4,5)-tetrakisphosphate
PTEN: phosphatase and tensin homolog
cryo-EM: cryo-electron microscopy
HDX-MS: hydrogen-deuterium exchange mass spectrometry
XL-MS: cross-linking mass spectrometry
SEC-SAXS: small-angle X-ray scattering
FL: full-length
RMSD: root mean squared deviation
COMs: centers of mass
SDS-PAGE: sodium dodecyl sulfate–polyacrylamide gel electrophoresis
mant-GTP: N-methyl-anthraniloyl-GTP
P(r): pair distance distribution
EOM: employed ensemble optimization method
WT: wild-type
MBP: maltose binding protein
GST: glutathione S-transferase
DTT: dithiothreitol
HEPES: 4-(2-hydroxyethyl)-1-piperazineethanesulfonic acid
NaCl: sodium chloride
EDTA: ethylenediaminetetraacetic acid
CV: column volumes
IPTG: isopropyl β-D-1-thiogalactopyranoside
MWCO: molecular weight cutoff
DDM: n-Dodecyl-Beta-Maltoside
PNCC: Pacific Northwest Center for Cryo-EM
BIS: beam image shift
CTF: contrast transfer function
FSC: fourier shell correlation
GuHCl: guanidine hydrochloride
LC/MS: liquid chromatography/mass spectrometry
D_2_O: deuterium oxide
Tris: Tris(hydroxymethyl)aminomethane
MgCl_2_: magnesium chloride
Rg: radius of gyration
Vc: volume of correlation

