## Supplemental Information for "P-Rex2 exhibits unique structural features and regulatory mechanisms distinct from the closely related RhoGEF P-Rex1"

**Contents**

Supporting Information Figures S1-S10

**Figure S1. P-Rex2 cross-linking mass spectrometry (XL-MS) data agree with the cryo-EM structure of P-Rex2**
**Figure S2. P-Rex2 cryo-EM data collection and processing**
**Figure S3. Overview of P-Rex2 cryo-EM data processing pathway**
**Figure S4. Interpretability of maps and fit of P-Rex2 model**
**Figure S5. 2D class averages show that $IP_4$ binding does not stabilize a PH–IP4P interaction to allow resolution of the P-Rex2 IP4P subdomain**
**Figure S6. P-Rex2 SEC-SAXS data analysis**
**Figure S7. BilboMD re-analysis of P-Rex1 DH/PH and DH/PH-DEP1 SEC-SAXS data**
**Figure S8. SDS-PAGE of P-Rex2 DH/PH-DEP1 and DH/PH proteins used in Figure 6 GEF activity assays**
**Figure S9. Comparison of the αH/αI interface between P-Rex1 and P-Rex2**
**Figure S10. HDX-MS data on P-Rex2 with and without $IP_4$**

A.

| P-Rex1 Homologous Residue 1 | P-Rex1 Homologous Residue 2 | Distance (Å) | P-Rex2 Residue 1 | P-Rex2 Residue 2 | Distance (Å) |
|---|---|---|---|---|---|
| 141 | 938 | 97.7 | 115 | 904 | Not modeled, unlikely to be within cross-linking distance |
| 154 | 1272 | 61.9 | 128 | 1212 | Unknown with current model |
| 241 | 1272 | 48.3 | 215 | 1212 | Unknown with current model |
| 164 | 1502 | 32.6 | 138 | 1442 | Not modeled, likely to be within cross-linking distance |
| 395 | 1502 | 22.2 | 364 | 1442 | 35.8 |
| 368 | 1502 | 32.8 | 337 | 1442 | 56.5 |
| 418 | 513 | 33.7 | 387 | 481 | 20.8 |
| 415 | 1502 | 42.0 | 384 | 1442 | 15.9 |
| 418 | 1502 | 45.1 | 387 | 1442 | 17.0 |
| 429 | 1502 | 54.4 | 397 | 1442 | 29.9 |

B.

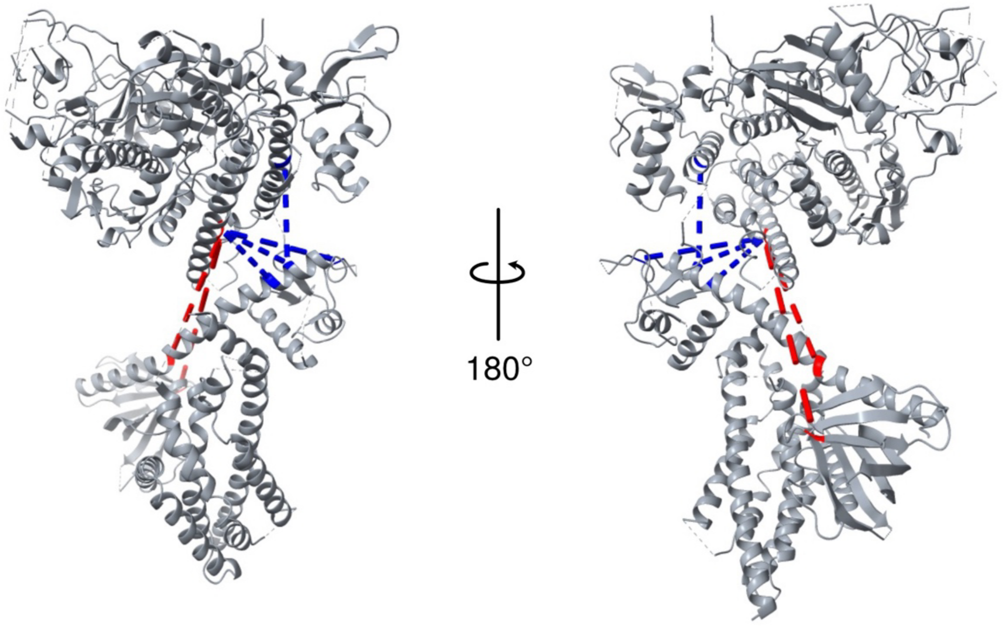


**Figure S1. P-Rex2 cross-linking mass spectrometry (XL-MS) data agree with the cryo-EM structure of P-Rex2.** A) Using P-Rex2 cross-linking mass spectrometry data from D'Andrea et al. (2021), we analyzed cross-links between residues in the N-terminal module and the C-terminal core. Looking at homologous residues in P-Rex1, we calculated the distances between Cα atoms of cross-linked residues using the structure of autoinhibited P-Rex1 (PDB: 8TUA). After obtaining the cryo-EM structure of P-Rex2, we performed the same calculations. B) P-Rex2 XL-MS data for cross-links between residues in the N-terminal module and the C-terminal core shown plotted onto the P-Rex2 structure. Blue dashes represent cross-linked residues that are <30 Å apart and red dashes represent cross-linked residues that are > 30 Å apart.

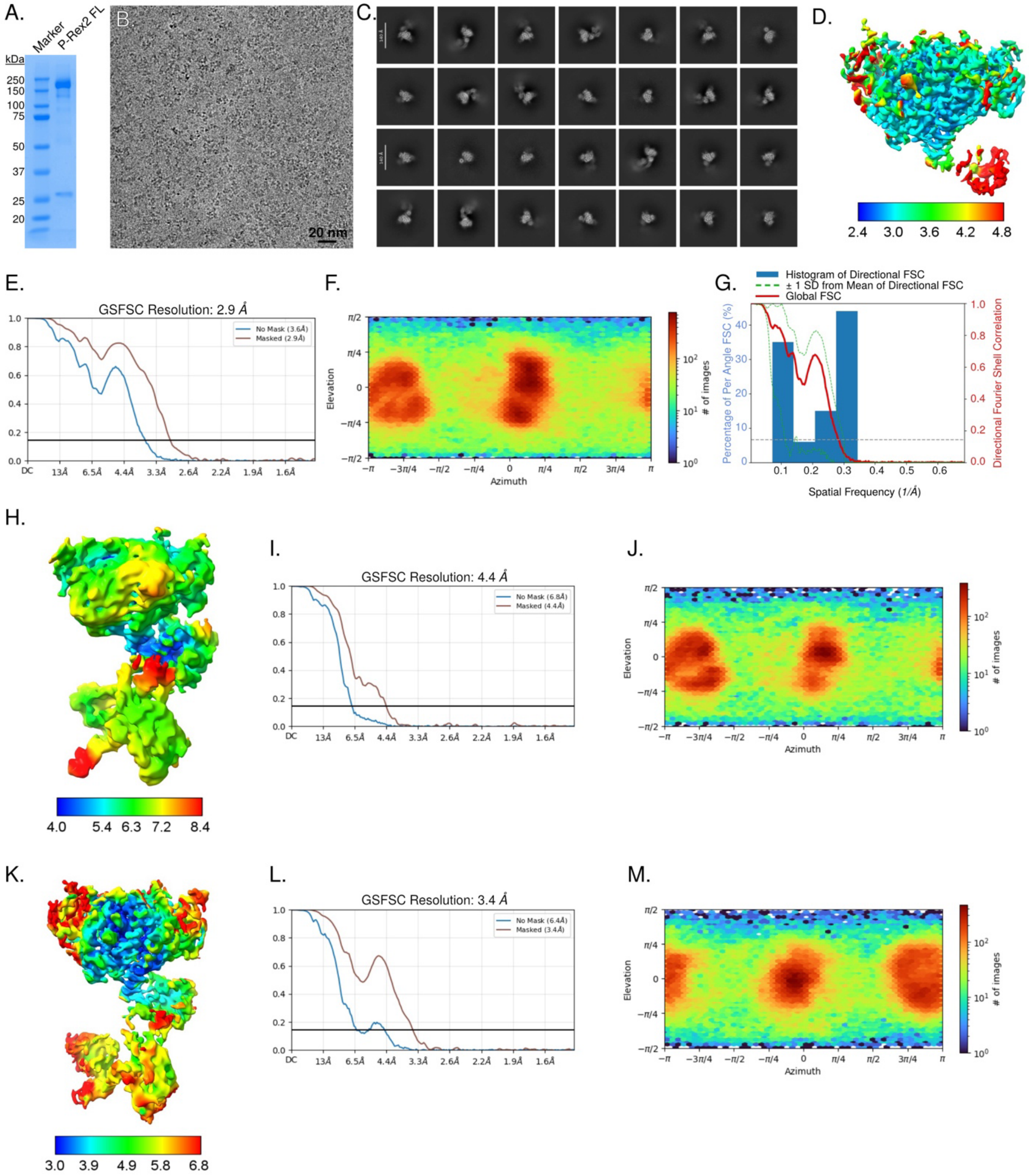


**Figure S2. P-Rex2 cryo-EM data collection and processing.** A) SDS-PAGE of purified P-Rex2 FL. B) Representative micrograph. C) Best 2D class averages from P-Rex2 data processing. D-F) Consensus map colored by local resolution and the corresponding FSC curve and viewing direction distribution plot. G) 3D FSC histogram and directional FSC plot calculated by the 3D FSC program (https://3dfsc.salk.edu/). Sphericity is reported to be 0.734 out of 1. Global resolution is reported as 3.6 Å. H-J) Map from local refinement of the N-terminal module colored by local resolution and the corresponding FSC curve and viewing direction distribution plot. K-M) Map from local refinement of the whole particle colored by local resolution and the corresponding FSC curve and viewing direction distribution plot.

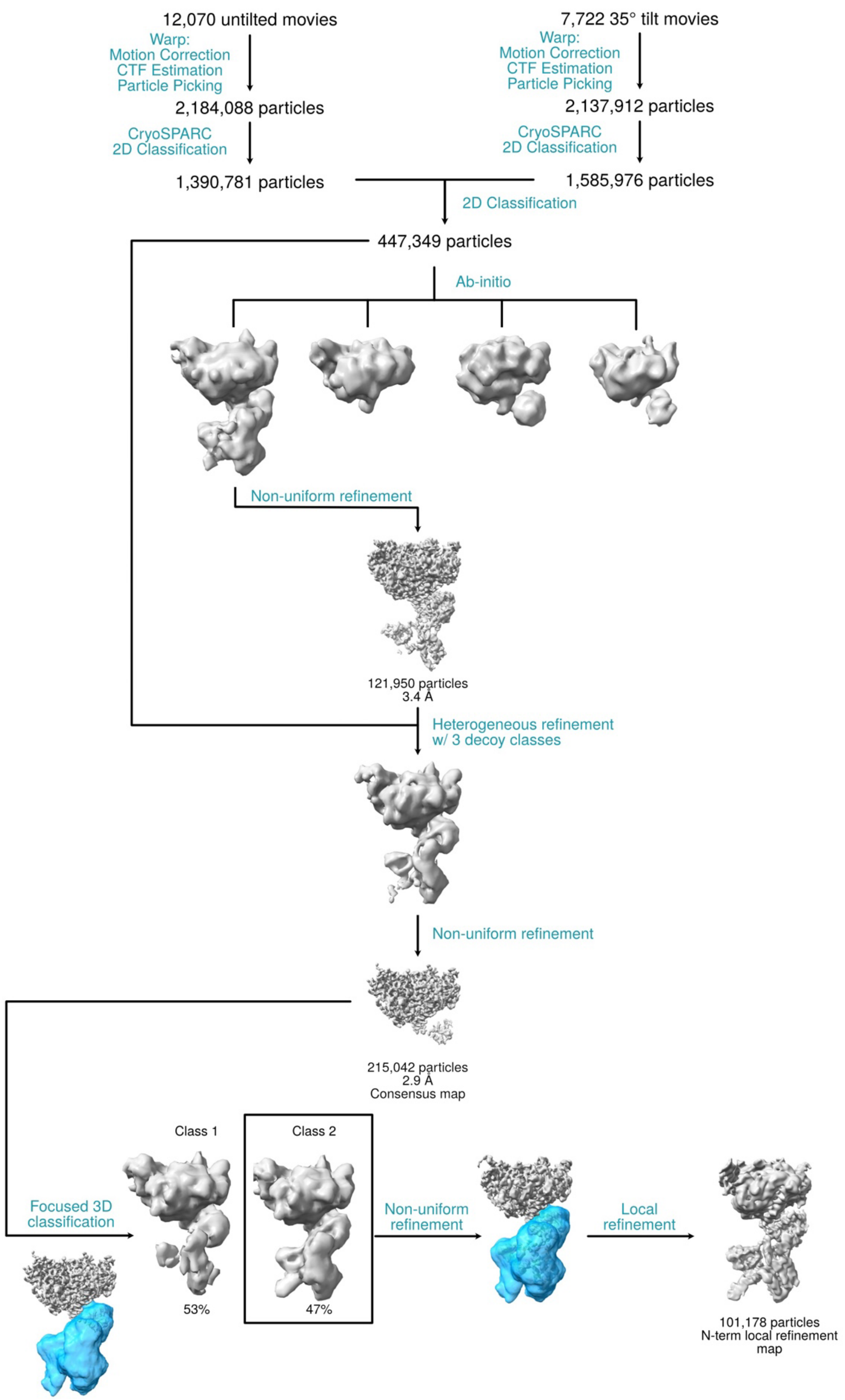


**Figure S3. Overview of P-Rex2 cryo-EM data processing pathway.**

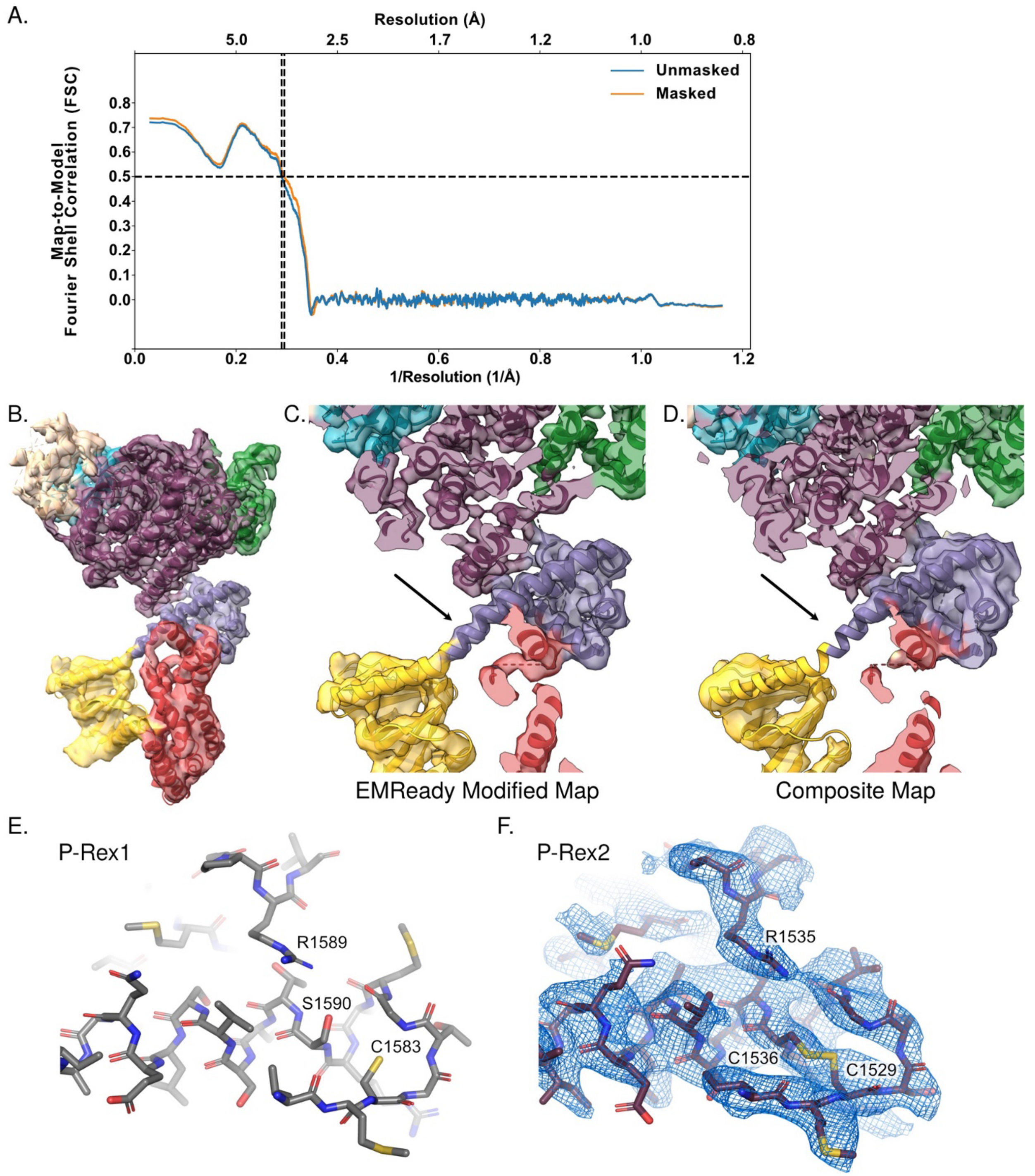


**Figure S4. Interpretability of maps and fit of P-Rex2 model.** A) Map-to-model FSC of the composite map. B) A density modified map generated with EMReady using the map from P-Rex2 whole particle local refinement. C-D) This map was especially useful in regions with poor resolution in the composite map such as in the helix bridging the PH and DEP1 domains (black arrow). E) P-Rex1 residues representing part of the catalytic triad of the proposed phosphatase site. F) P-Rex2 residues of the homologous phosphatase site and composite map density representing them, showing that this area is well-resolved in the map.

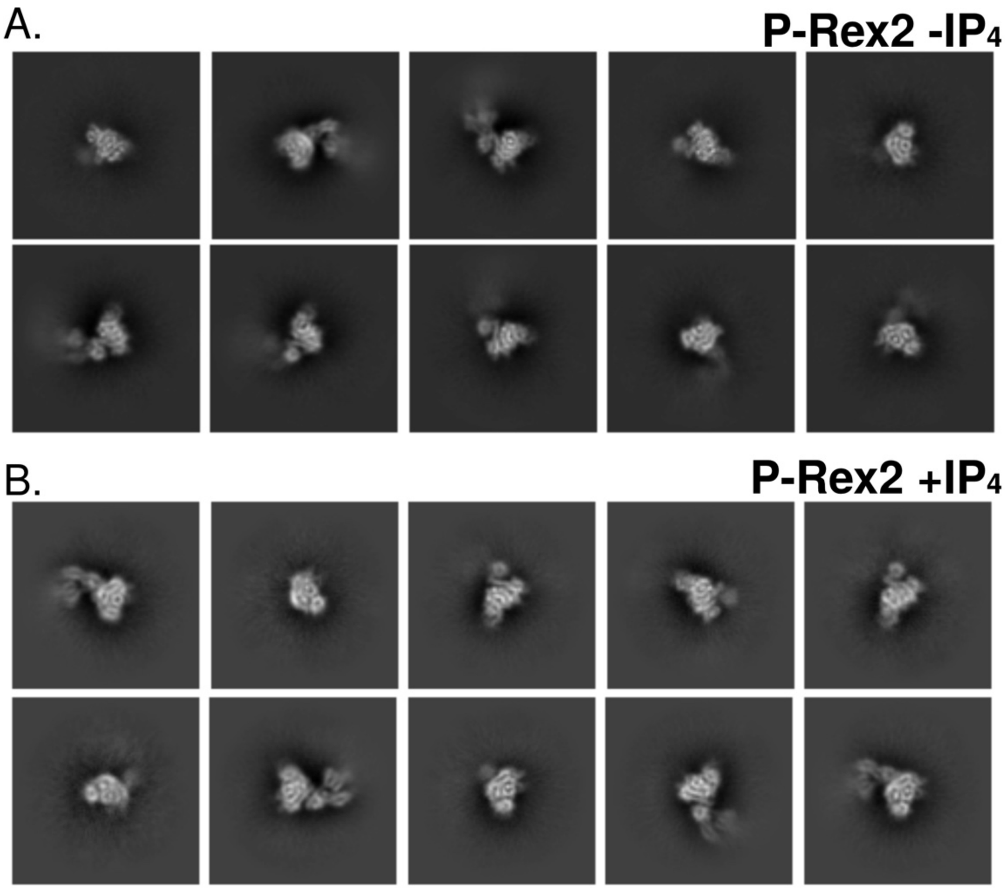


**Figure S5. 2D class averages show that $IP_4$ binding does not stabilize a PH–IP4P interaction to allow resolution of the P-Rex2 IP4P subdomain.** Two small datasets were collected on a Glacios on P-Rex2 samples in the A) absence and B) presence of $IP_4$. Datasets were processed in the same manner.

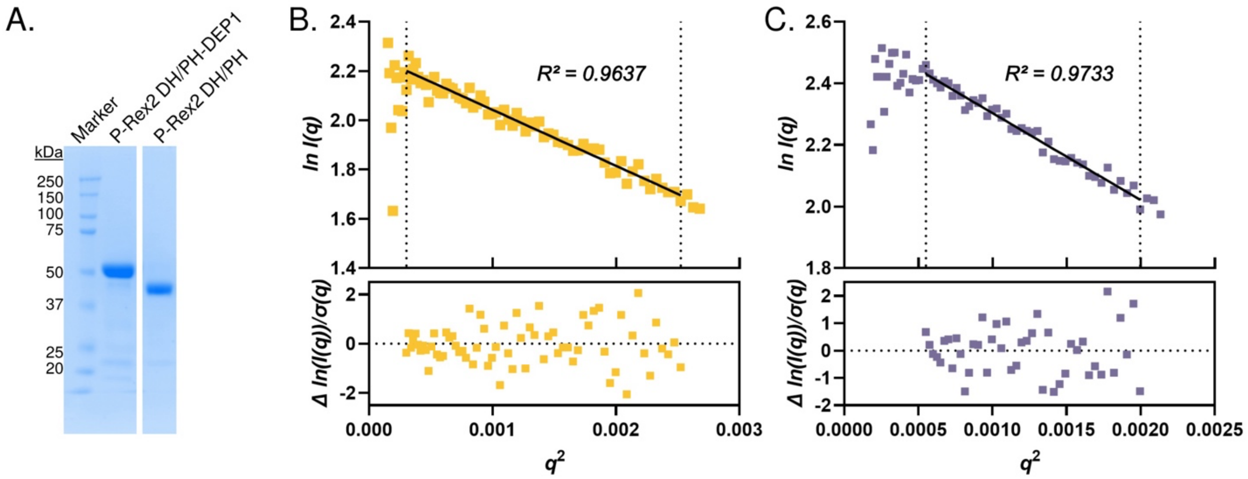


**Figure S6. P-Rex2 SEC-SAXS data analysis.** A) SDS-PAGE of purified P-Rex2 DH/PH and DH/PH-DEP1 constructs used in SEC-SAXS and GEF activity assay experiments. B) P-Rex2 DH/PH and C) P-Rex2 DH/PH-DEP1 Guinier plots.

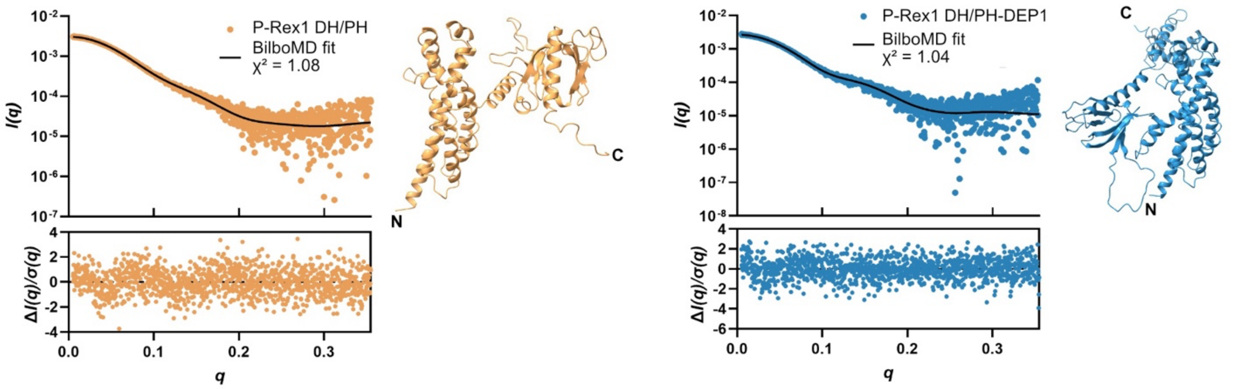


**Figure S7. BilboMD re-analysis of P-Rex1 DH/PH and DH/PH-DEP1 SEC-SAXS data.** Published SEC-SAXS datasets of P-Rex1 DH/PH and DH/PH-DEP1 (SASDHY9 and SASDHW9; Ravala et al., 2020) were analyzed using the BilboMD program. Scattering intensity plots of P-Rex1 DH/PH (orange) and DH/PH-DEP1 (blue) fit with the BilboMD model. The output BilboMD model is shown to the right of the plot. Normalized fit residuals are shown in the bottom panel.

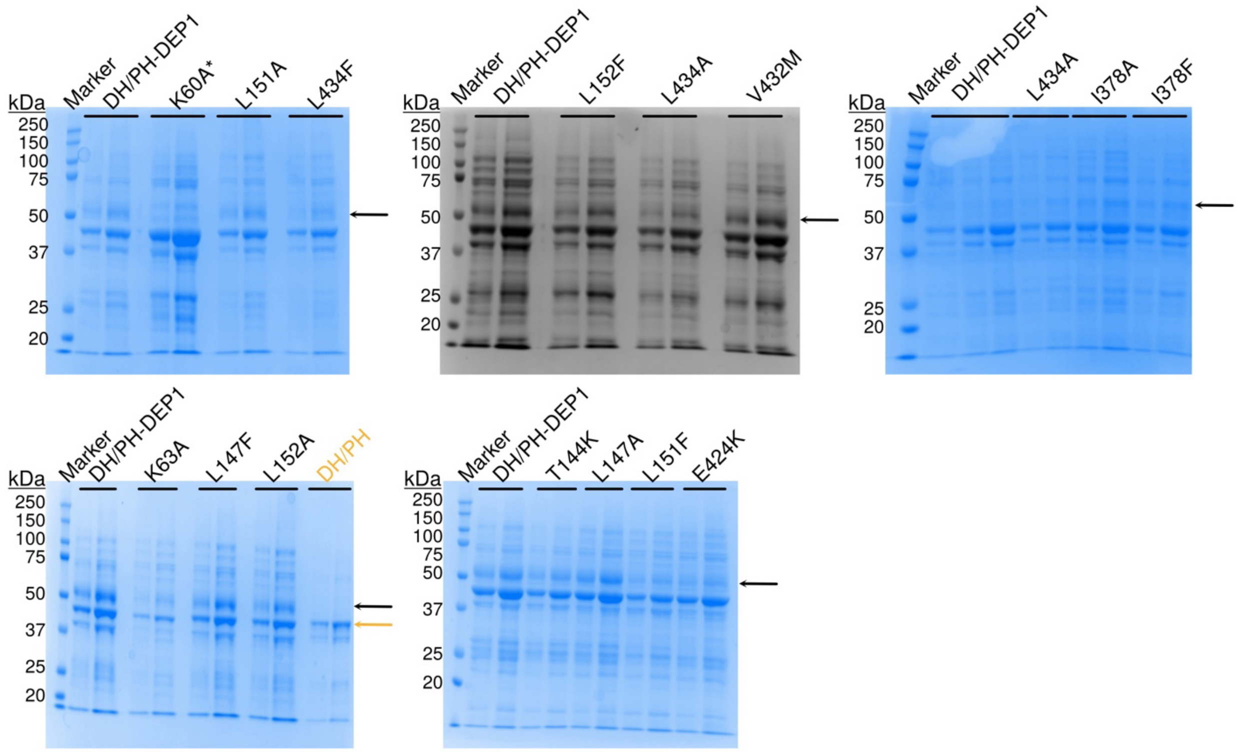


**Figure S8. SDS-PAGE of P-Rex2 DH/PH-DEP1 and DH/PH proteins used in Figure 6 GEF activity assays.** The black arrow indicates the band that represents each P-Rex2 DH/PH-DEP1 construct (~51 kDa). The yellow arrow indicates the band that represents P-Rex2 DH/PH (~44 kDa). For each construct, two lanes were run with different sample volumes to aid in estimating sample concentration.

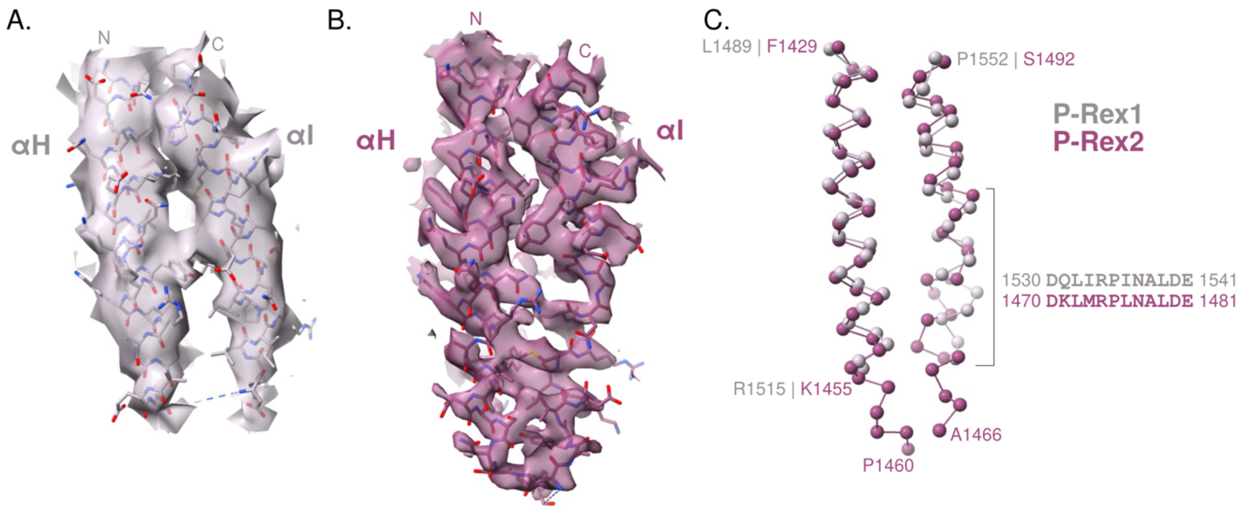


**Figure S9. Comparison of the αH/αI interface between P-Rex1 and P-Rex2.** Cryo-EM maps with the models fit showing how the regions around the αH and αI helices in A) P-Rex1 (PDB: 8TUA, EMDB-41621) and B) P-Rex2 are significantly different. There is also greater resolvability in the loop connecting these features in P-Rex2. C) The isolated Cα trace shows that this difference is most pronounced in the alternate structure from D1470 to E1481 in P-Rex2.

**Figure S10. HDX-MS data on P-Rex2 with and without $IP_4$.** Ribbon maps representing HDX-MS experiments with P-Rex2 alone and bound to $IP_4$. Also shown are the changes in exchange rates that occur in P-Rex2 upon $IP_4$ binding.

### Ribbon Map of P-Rex2 (% deuteration)

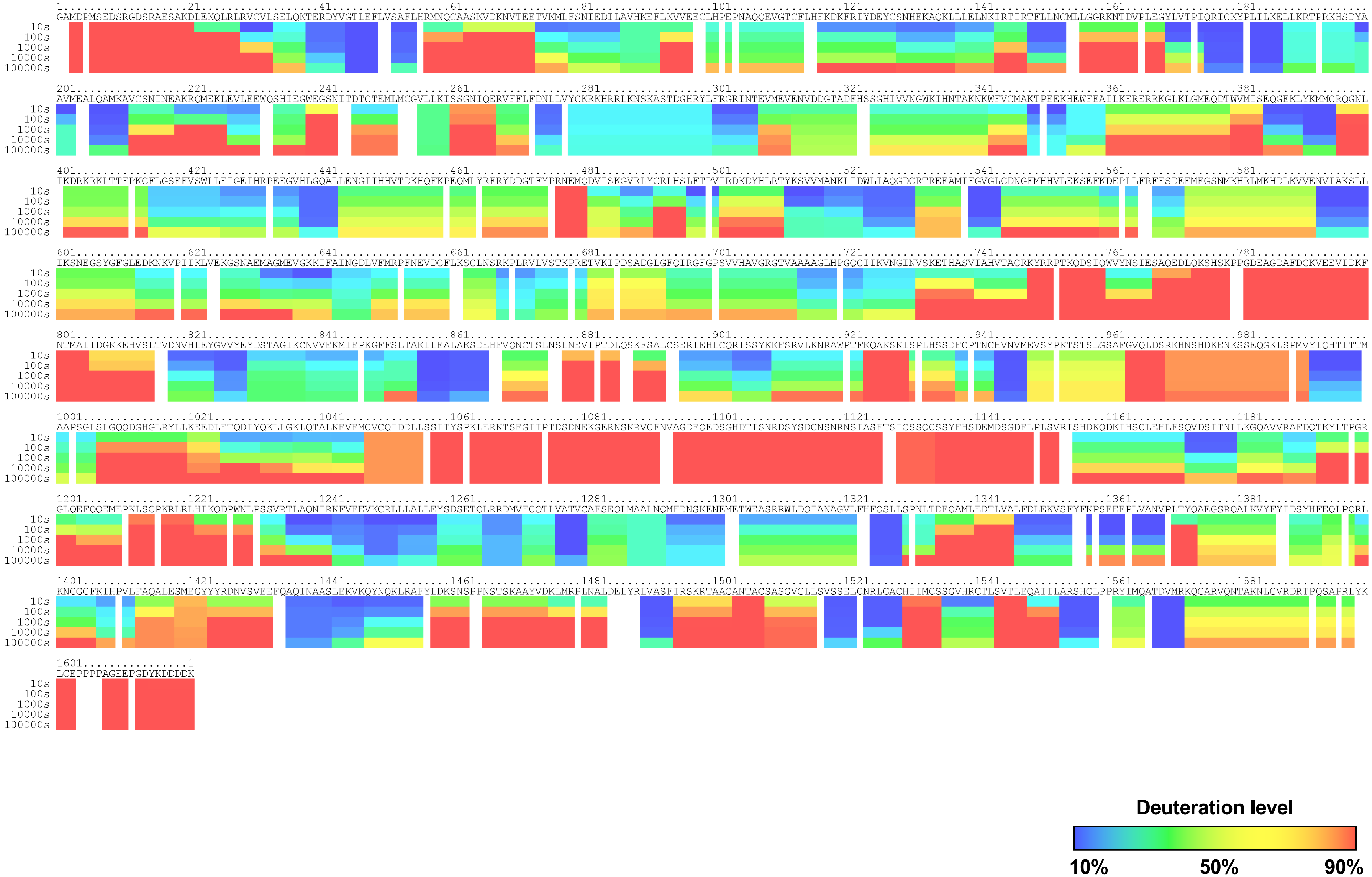

### Ribbon Map of P-Rex2 in P-Rex2•$IP_4$ Complex (% deuteration)

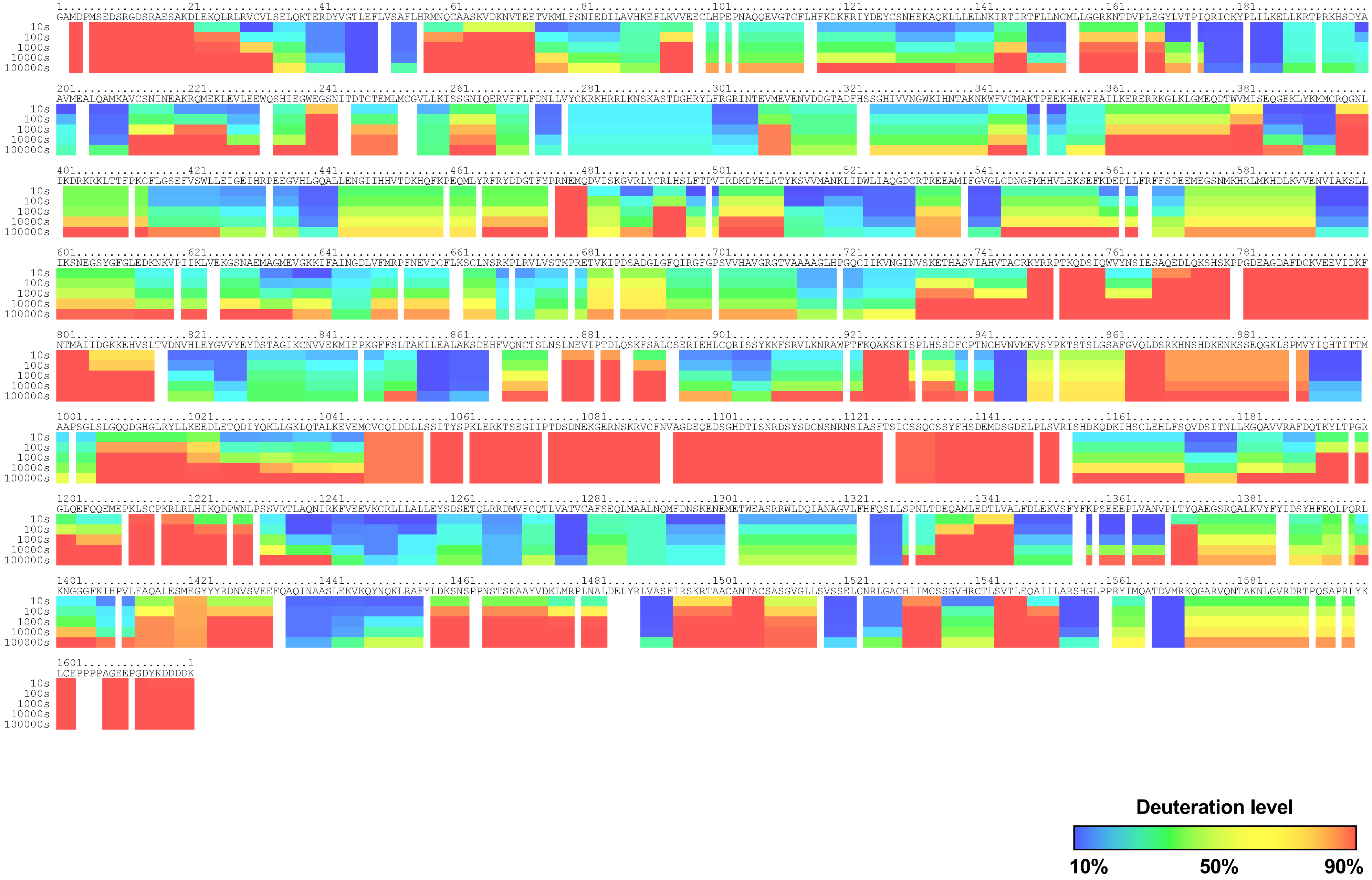

### Influence of $IP_4$ on Exchange in P-Rex2 (% deuteration)

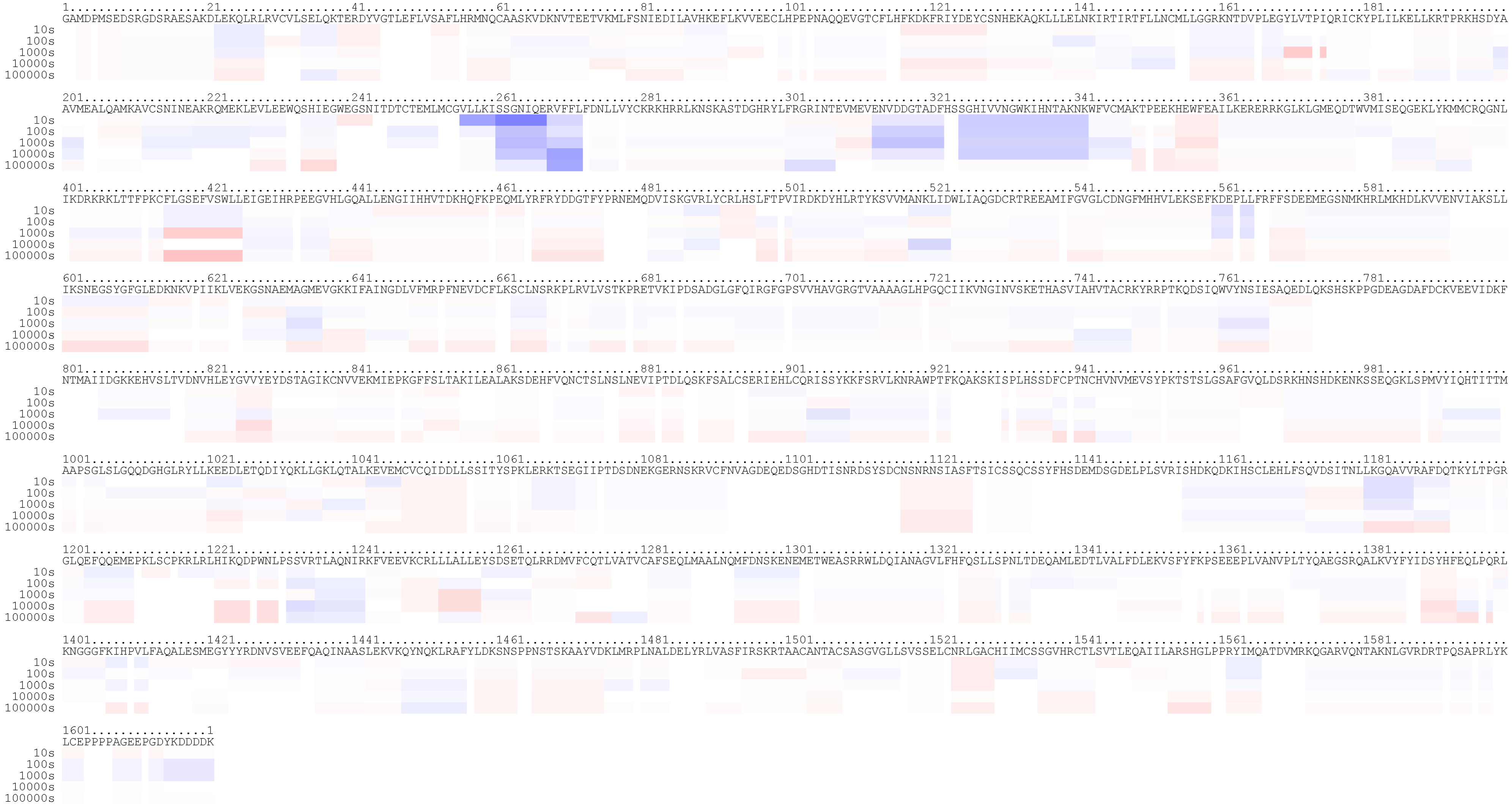


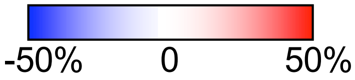


Blue indicates regions that exchange slower in the presence of $IP_4$.
Red indicates regions that exchange faster.
